# An integrative analysis of the biological clock hypothesis in human gut microbiome

**DOI:** 10.1101/2022.03.22.485238

**Authors:** Anh D. Moss, Shan Sun, Ivory C. Blakley, Anthony A. Fodor

## Abstract

**Background:** While previous studies have explored the relationship between aging and the gut microbiome, it remains unclear how consistent and reproducible this association is across different cultures and groups. We performed an integrative analysis with 11 independent datasets from nine different countries to test the idea that the aging gut microbiome can be viewed as a biological clock, in which microbial changes associated with age are consistent and measurable across distinct datasets.

**Results:** As expected, our Principal Coordinate Analysis found strong batch effects with study ID by far the strongest signal across datasets. Despite this large batch effect, we found a consistent signal across studies that was largely driven by sample size with only our larger cohorts showing taxa in common associated with age. Likewise, Shannon diversity and richness were not consistently associated with age across the 11 datasets, but some positive correlations with richness and host age were observed among the four largest cohorts. The taxon with the most potential as a biomarker for the aging human gut microbiome is genus *Bifidobacterium*, with significantly negative associations with host age in three out of the four datasets that had more than 200 samples.

**Conclusion:** The driving force behind low reproducibility of association of age with the microbiome in previous studies appears to be inadequate sample size rather than structural differences in the microbial community based on cohort characteristics. Results from a power analysis suggest that future studies on the aging human gut microbiome consider on the order of 100-300 samples to consistently observe an age signal. With these larger sample sizes, parametric and non-parametric model yield broadly similar power.

## Introduction

A fundamental question in biology is why we age and how to precisely define aging within biological parameters. The rate of living theory, based on observations that species with higher metabolism rates often have shorter lifespans, postulates that metabolism rate and lifespan are correlated (1, 2). The free radical theory of aging expands upon the rate of living theory and suggests that oxidative stress, caused by accumulated reactive oxidative species (ROS) from metabolic processes, lead to cellular damage and decreased physiological function (3). Additional studies support oxidative stress as a physiological mechanism of aging (4, 5). Building on the concept of aging as a result of accumulated cellular damage, some studies suggest that reactive oxidative species also potentially damage telomeres, ultimately decreasing telomere length and increasing risk of cellular mortality (6–8).

The human gut microbiome has been implicated in many diseases. Developmental changes of the gut microbiome composition have been reported across all age groups from infants to the elderly (9–23). The composition of the newborn human microbiome is largely determined by environmental exposure, such as delivery mode, and within the first six months to two years of life, the infant gut microbiome becomes more like the adult microbiome (9, 10, 12). At the other extreme of life, many studies have asked if age-related microbiome changes increase the risk of disease development (11, 13–23). Some studies have suggested that gut microbiome biodiversity is generally stable throughout adulthood (14, 16, 24). While these studies have produced valuable insights into how the microbiome changes with age, it remains an open question if aging causes the same changes in the gut microbiome across different cohorts and cultures and could therefore be considered as a biomarker of risks associated with aging.

In this study, we performed an integrative analysis across 11 previously published publicly available datasets in which we looked at associations with age. Within each dataset, we used Principal Coordinate Analysis (PCoA) and Kendall correlation tests to ask how age is associated with the microbial community. We use “p-value vs p-value” plots (Fig. 1) to visualize p-value trends between a given pair of cohorts, from which we interpret the resulting correlation as reproducibility of the age signal. The direction of the association in these plots is captured by multiplying the log10 p-value by −1 if the Kendall correlation estimate was positive. The red quadrants in these plots therefore indicate the regions of significant positive (upper right) and negative (lower left) associations between age and microbial shifts in both cohorts (Figure 1). The leftmost panel of Fig. 1 reflects a null hypothesis of no age signal. A lack of age signal would result in either a tight cluster of p-values around the center (origin) of the plot because neither cohort has taxa significantly associated with host age, or as a loosely clustered group of p-values, in which a few taxa might be correlated with age in one or both cohorts, but no more than might be expected from random false positives. Inconsistent signals between cohorts can be seen as a negative correlation with p-values falling into the black quadrants (upper left and lower right), where a taxon is significantly increased in one cohort but significantly decreased in another, or as a dispersion of p-values that reflect a particular pattern in one cohort but not the other. The alternative hypothesis, expressed in the middle and right panels, states that an age signal exists in the human gut microbiome and is consistent across some (Figure 1, middle panel) or all (Figure 1, right panel) cohorts. In the rightmost panels, we see a model of the aging gut microbiome following the clock hypothesis, in which the effect size of host age on microbial abundance is strong enough to be observed across multiple different independent cohorts.

**Figure 1.**
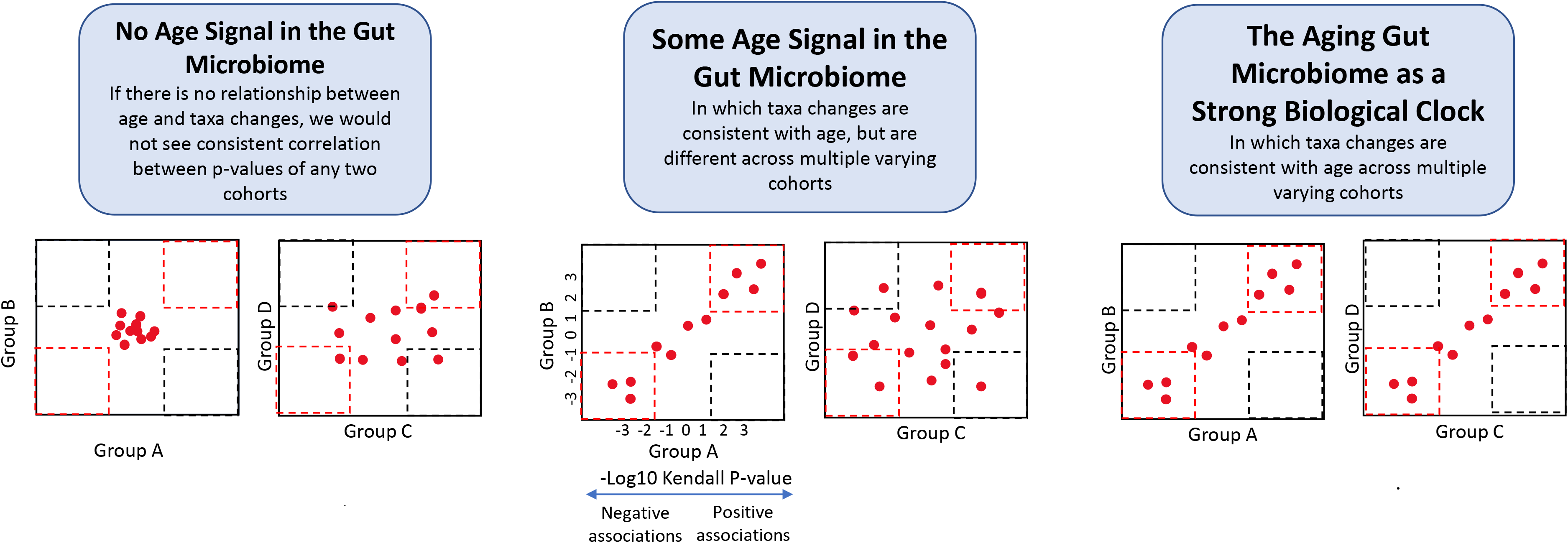
A visual summary of how different conceptual models of the aging gut microbiome would impact p-value vs p-value plots, a central visualization in our study (see text in introduction). In these plots, we compared unadjusted log_10_ Kendall p-values for shared taxa between two cohorts. Groups A-D represent theoretical example of independent datasets which could be entirely independent in their associations with age (left panel), show consistent patterns of associations with age for same cohorts but not others (middle panel) or show consistent associations with age across all cohorts (right panel).

The results from our study favors the middle panel in which consistent changes are seen in some but not all cohorts. We argue, however, that the differences between studies may be driven by the small sample size of some studies and the overall small effect size of associations with age. We present power analyses that suggest a minimal sample size of around 100-300 subjects to reliably see associations with age by either parametric or non-parametric tests.

## Methods

### 16S rRNA Gene Sequence Datasets of the Human Gut Microbiome

For data collection, we performed a literature search on Google Scholar for publicly available 16S rRNA gene sequence data of the human gut microbiome with accessible metadata of host ages. We included only datasets with an age range spanning across more than two decades (Supp. Figure 1). The datasets were downloaded from the National Center for Biotechnology Information Sequence Read Archive (NCBI SRA), as well as most of their respective metadata, although some metadata were obtained from individual websites (Table 1). We found a total of 11 cohorts from nine different countries, including Canada, the United States, the United Kingdom, China, Colombia, Spain, Sweden, France, and Germany (Table 1). These datasets are based on studies of five different health conditions, including HIV (25), colorectal cancer (26, 27), irritable bowel disease (28), obesity (29, 30), and Type 2 diabetes (29, 30). Four datasets consist of only healthy participants (16, 27, 31, 32). The sequence data are based on human stool samples, except for data from the Morgan and Zeller Germany cohorts. The Morgan cohort had stool (n=134) and biopsy tissue (n=94). We ran the stool and biopsy samples separately and saw no significant results for either tissue type (Supp. Figure 4B). When we combined these samples together (n=228), we saw a single taxon significantly associated (*Bifidobacterium*) giving some further support to the idea that *Bifidobacterium* is consistently associated with age in large sample sizes. In this manuscript, we chose to report only the stool samples to ensure the most compatibility with the majority of other cohorts and did not pursue the merged cohort, in part because it was unclear if some of the samples were obtained from the same individuals which would put our models in violation of assumptions of independence.

**Table 1.**
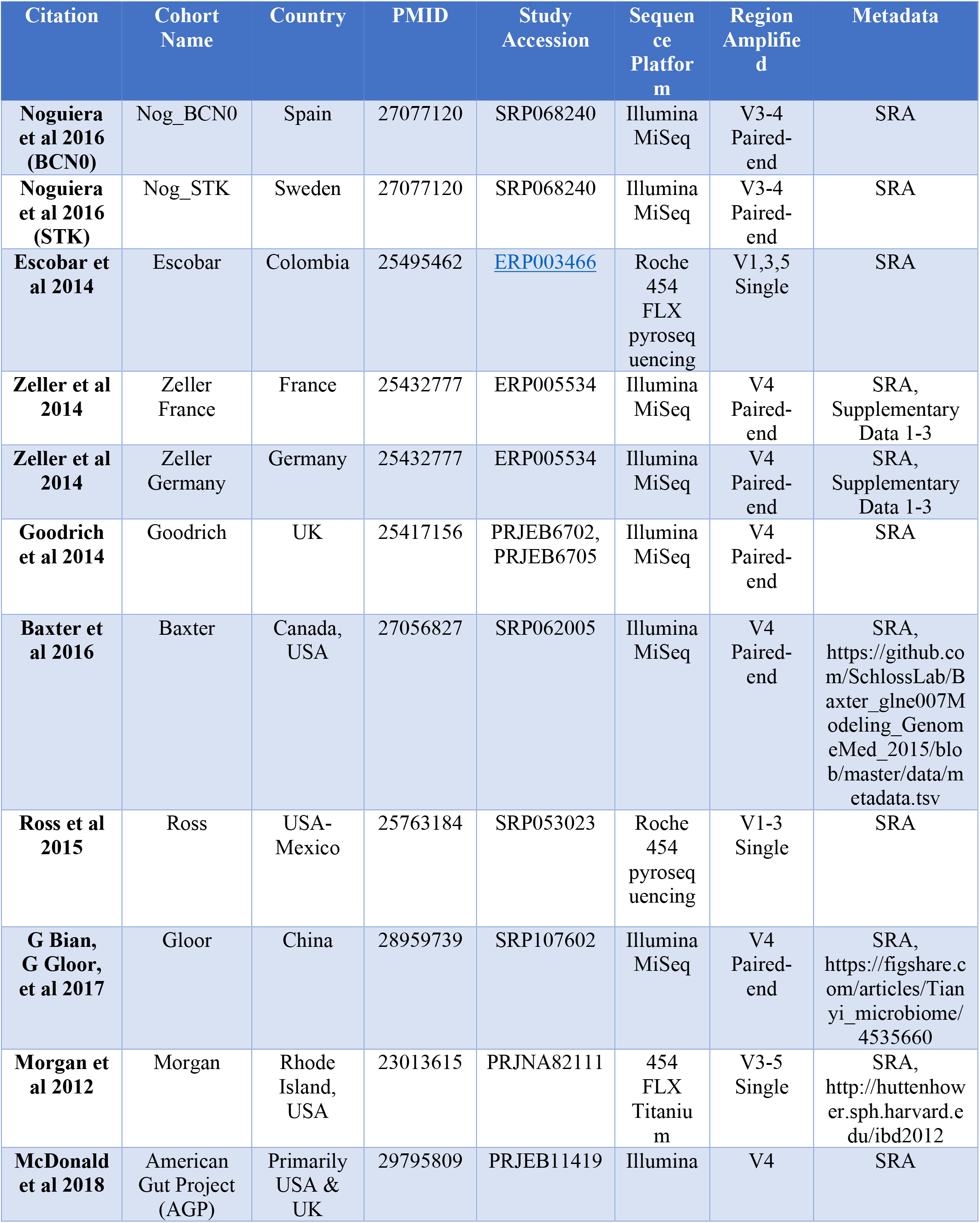
Summary of the 11 datasets in this integrative analysis. The second column, “Cohort Name”, specifies the cohort names used in our analysis respective to each individual study.

In the Zeller Germany cohort, matched samples of colonic tissue of patients with colorectal cancer were collected, one from a tumor and one from a nearby morphologically healthy tissue (27). For this study, we included sequences based only on the healthy samples (removing the matched tumor samples from our analysis), and all analyses were performed at the genus level.

We excluded samples without host age metadata or had duplicates from the same host. In the Gloor dataset, stool samples were collected from two different cohorts: from the general population and from police and military training populations, which consisted mostly of young adult males (16). There were no available metadata for the military training cohort, so those samples were excluded from our integrative analysis. Out of the 1,352 samples from the general population cohort, we matched 803 samples with metadata from the NCBI SRA database so we consider the Gloor sample size to be 803.

Since only a few cohorts in this analysis had longitudinal sampling, to prevent sampling bias and remain consistent with the other datasets, we excluded the longitudinal samples. The available data from the Goodrich cohort consisted of samples from 951 individuals, of which 89 were part of a longitudinal cohort that provided multiple fecal samples over time. As such, we included 835 unique samples collected at only one time point from the Goodrich cohort (31).

### Pre-Processing Sequence Data

For quality control, we used FastQC (v. 0.11.5) to assess sequence quality, then used Trimmomatic (v. 0.38) to remove adapters and low-quality sequences (33, 34). In the Trimmomatic step, we used the following parameters: SLIDINGWINDOW 4:15, HEAD:20 to remove the first 20 base pairs, a minimum sequence length of 50, and all sequences cropped at 180 base pairs long. For the AGP cohort, we used DADA2 instead of Trimmomatic to simply set the maximum sequence length to 120 base pairs because this dataset had shorter sequence read lengths. However, all the other samples that were processed with Trimmomatic were also ran through DADA2 with the same respective parameters (maximum sequence length of 180 bp). For datasets with paired-end sequences, we used only the forward reads. We used QIIME2 (v.2021.2) with the SILVA database to taxonomically profile the 16S rRNA gene sequence data, from which we used counts at the genus level for the downstream analysis (35). Then, we normalized the genus abundance with the following equation to correct for different sequencing depth across samples:

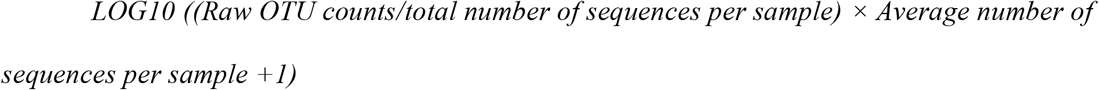

This equation is designed to minimize differences from adding the pseudo-count due to read depth. We performed this normalization schema at the individual cohort level, as well as subgroup and pooled samples level of analysis.

### Statistical Analysis

We used the vegan package in R (v.3.6.0) for Principal Coordinate Analysis (PCoA) based on the Bray-Curtis distance with the *capscale* function, PERMANOVA tests using the *adonis* function, and alpha diversity analysis (Shannon Diversity Index and richness) with the *diversity* function and *specnumber* function, respectively.

We used the base-R stats package for inferential statistical tests (*cor.test*, *wilcox.test, t.test,* and *lm* functions), with age as the independent variable and genus abundance as the dependent variable for each dataset.

## Results

We performed an integrative analysis across multiple cohorts of human gut microbiome datasets from North America (26, 28, 30, 32), China (16), South America (29), and Europe (25, 27, 31, 32) (Table 1). To account for the varying age ranges of each dataset, we designated the following subgroups with similar age ranges as “Younger Range”, “Middle Range”, and “Full Range”. The Younger Range group includes the three cohorts (Noguiera (BCN0), Noguiera (STK), and Escobar) that range from 19 to 78 years with a cumulative average of 42 ± 11 years (Supp. Figure 1, top panel). The Middle Range group has primarily middle age and elderly subjects (average= 61 ± 11 years, range from 23 to 96 years); this group includes the Zeller (France), Zeller (Germany), Goodrich, Baxter, and Ross cohorts. Lastly, the Full Range subgroup includes the remaining three cohorts (Gloor, Morgan, and AGP) that have the broadest age range and did not fit into the other two groups, with an average host age of 45 ± 26 years and range of 3 to 109 years (Table 2).

**Table 2.** Demographic summary of each dataset.

| Cohort Name | Health Condition | Sample Size | Age Range<br>(Mean ± SD<br>Years) |
| --- | --- | --- | --- |
| **Nog_BCN0** | HIV | 156 | 43 ± 10 |
| **Nog_STK** | HIV | 84 | 41 ± 12 |
| **Escobar** | Obesity | 30 | 38 ± 11 |
| **Zeller France** | Colorectal Cancer | 129 | 63 ± 9 |
| **Zeller Germany** | Colorectal Cancer | 48 | 65 ± 13 |
| **Goodrich** | Healthy | 835 | 61 ± 10 |
| **Baxter** | Colorectal Cancer | 490 | 60 ± 12 |
| **Ross** | Obesity | 63 | 53 ± 11 |
| **Gloor** | Healthy | 803 | 43 ± 36 |
| **Morgan** | Irritable Bowel<br>Disease | 134 | 35 ± 19 |
| **AGP** | Healthy | 1,368 | 47 ± 18 |
| **Totals** | -- | 4,140 | 51 ± 22 |

### Batch effect by cohort drives separation of variation across samples

To characterize the relationship between microbial composition and host age, we first performed a Principal Coordinate Analysis (PCoA) using the Bray-Curtis dissimilarity metric at the genus level in three ways: (i) for all samples pooled together across all datasets, (ii) with separate PCoAs for our three subgroups, and (iii) for each dataset separately. In our PCoA with all samples, we pooled 4,140 samples from all 11 cohorts and saw that the largest driver of difference in the PCoA plot was batch effect by study (Figure 2A). For pooled samples, we ran a PERMANOVA model with both cohort and age as terms. While both terms were significant for all permutations with p-value = 0.001, cohort (R^2^ = 0.223) explained much more of the variance in the data than age (R^2^= 0.00665).

**Figure 2.**
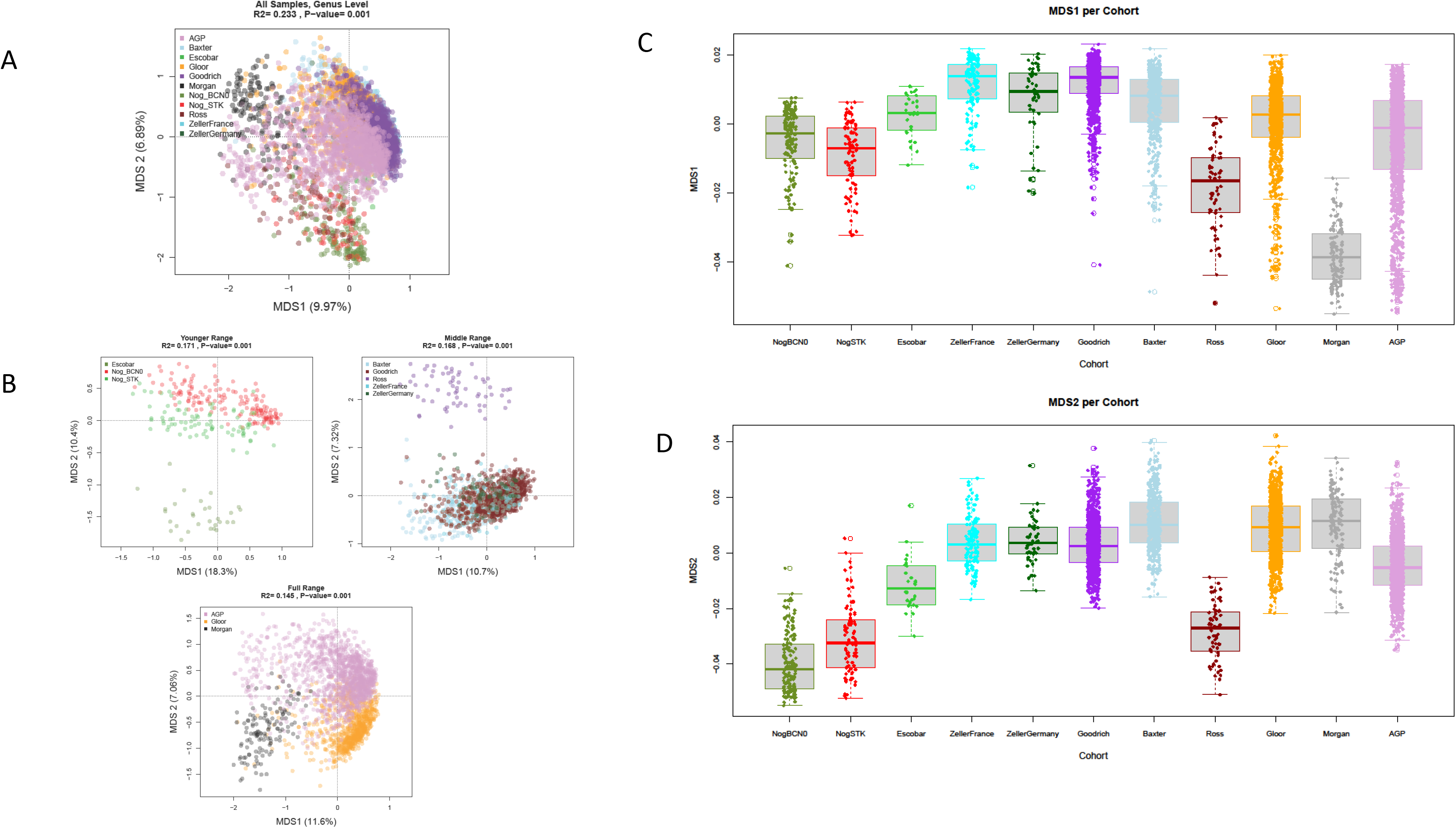
(A) PCoA plot for pooled samples, with points colored by study. The R2 values and p-values are from PERMANOVA tests with age as a continuous variable. (B) PCoA plots of genus abundance for samples at the subgroup level, with samples colored by study for the Younger Range, the Middle Range, and the Full Range. (C and D) A boxplot view of the first two MDS axes from the PCoA of pooled samples colored by cohort.

To see if datasets with similar age ranges still showed separation by batch effect, we performed PCoAs at the subgroup level with the Younger, Middle, and Full Range subgroups. We still observed separation by study across all three subgroups (Figure 2B).

Next, we performed PCoAs to characterize variations within each cohort to see if any variations were associated with host age. When we performed PERMANOVA tests for these PCoAs, seven of the 11 cohorts had significant associations between host age and the gut microbiome at a p-value threshold of 0.05 (Table 3). Six of these seven cohorts with significant PERMANOVA p-values had the largest sample sizes among the 11 datasets in this integrative analysis (Table 2 and 3). However, there was no trend between R^2^ values and sample size (Table 3).

**Table 3.** PERMANOVA p-value and R^2^ values for each individual cohort.

| Cohort | P-Value | R<sup>2</sup> Value | Significant Taxa | Age Range |
| --- | --- | --- | --- | --- |
| **NogBCN0** | 0.001 | 0.0462 | 32 | 49 |
| **NogSTK** | 0.098 | 0.0186 | 0 | 59 |
| **Escobar** | 0.013 | 0.0640 | 0 | 39 |
| **ZellerFrance** | 0.171 | 0.0099 | 0 | 52 |
| **ZellerGermany** | 0.769 | 0.0163 | 0 | 61 |
| **Goodrich** | 0.001 | 0.0042 | 25 | 63 |
| **Baxter** | 0.001 | 0.0079 | 10 | 60 |
| **Ross** | 0.355 | 0.0170 | 0 | 48 |
| **Gloor** | 0.001 | 0.0420 | 217 | 106 |
| **Morgan** | 0.029 | 0.0136 | 0 | 76 |
| **AGP** | 0.001 | 0.0076 | 75 | 86 |

In addition, we compared host age within each cohort along the first two MDS axes and assessed the correlation with the Kendall correlation test. In the first MDS axis, six cohorts had significant age-related variation with a significance threshold at 0.05 (Supp. Figure 2A). Within the second MDS axis, five cohorts had a significant association with host age (Noguiera BCNO, Baxter, Gloor, AGP, and Morgan) (Supp. Figure 2B).

While there were some significant age-related differences between the samples, most cohorts had a low PERMANOVA R^2^ value. The following three cohorts presented the highest R^2^ values: the Escobar (R^2^= 0. 0640), Noguiera_BCN0 (R^2^=0. 0462), and Gloor cohort (R^2^=0. 0420) (Table 3).

### There is little evidence of a consistent loss of diversity associated with age across the 11 datasets

To ask if alpha diversity in the human gut microbiome is associated with age, we calculated Shannon’s diversity for each of the 11 cohorts in our integrative analysis. Previous literature has suggested an interaction between Shannon diversity and sample depth (36). However, Shannon diversity calculated on both rarified and unrarefied samples were highly concordant (Supp. Figure 3). Therefore, we do not think that our Shannon diversity results are confounded by sequencing depth. We tested the associations of alpha diversity and richness against age with the Kendall correlation test and a p-value cutoff at 0.05. Across the 11 datasets, the Kendall tau correlations (Shannon diversity ~ age) ranged from −0.188 to 0.0679, with only the Goodrich cohort with a significant correlation between diversity and age at an uncorrected p-value < 0.05 (Kendall tau = 0.0679) (Figure 3A).

**Figure 3.**
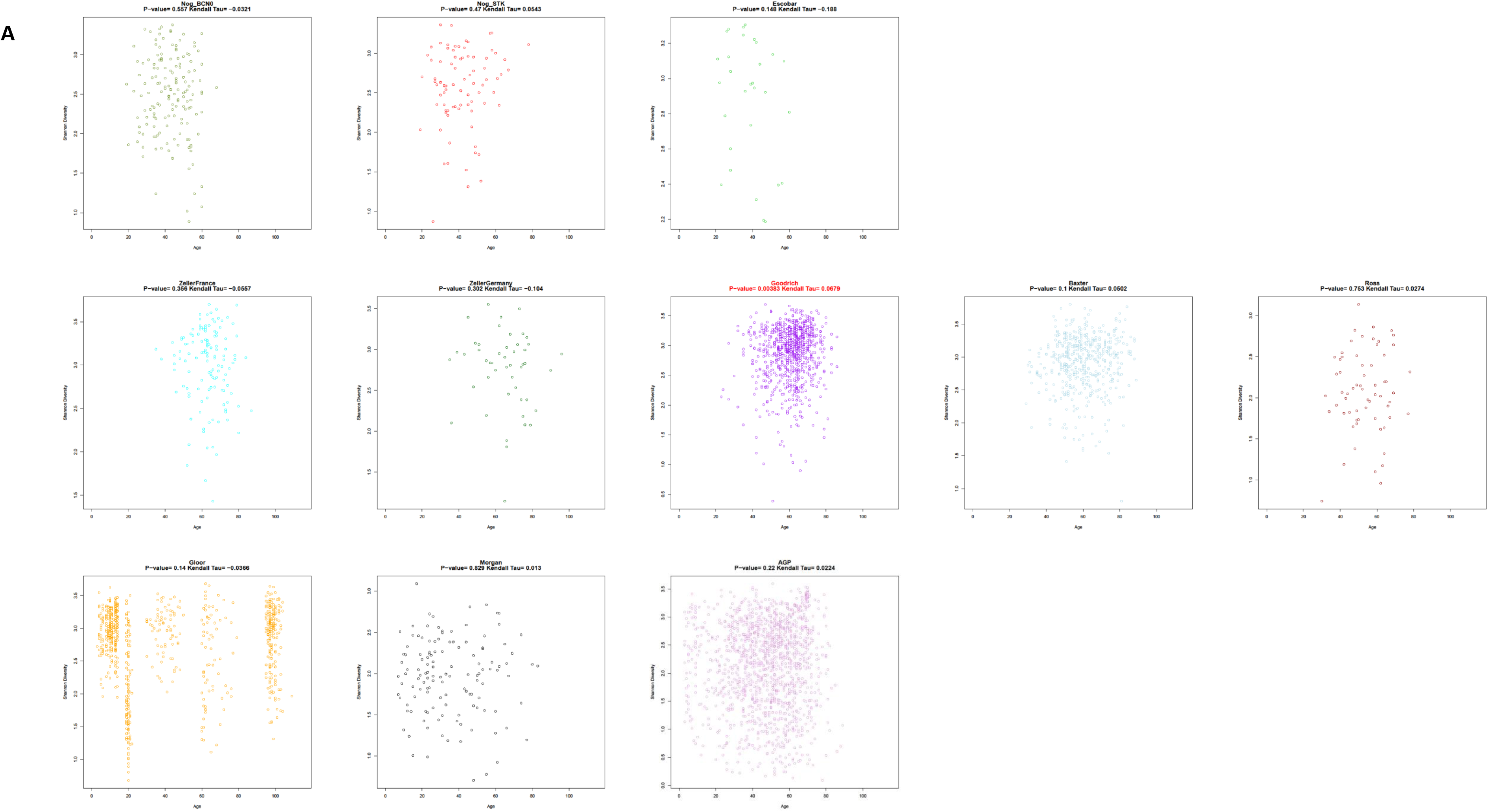
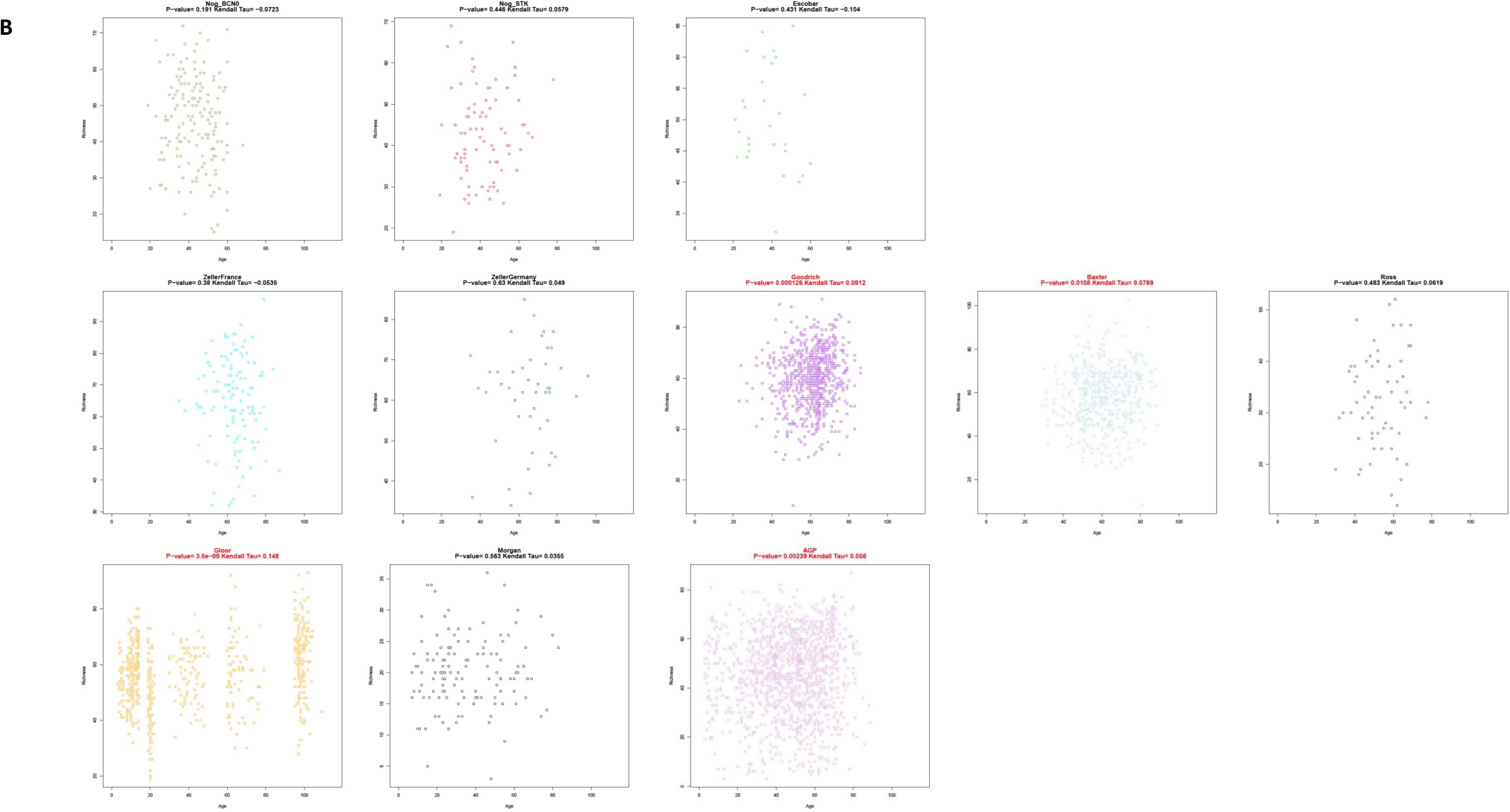
Scatter plots of (A) Shannon diversity and (B) richness for each of the ten datasets at genus level, with age as a continuous variable. Cohorts with significant correlations are highlighted in red text.

Taxa richness was significantly associated with age in four out of the 11 cohorts, all of which were the largest datasets in this study (Goodrich, Baxter, Gloor, and AGP) (Figure 3B). Among these four cohorts, we observed increased taxa richness with host age (Figure 3B). The Gloor cohort had the largest effect size (Kendall tau=0.148), followed by the Goodrich cohort (Kendall tau=0.0912) (Figure 3B).

### No specific taxa are significantly associated with host age across all cohorts

Since an analysis of MDS axes and diversity do not provide details on specific taxa, we next used the Kendall correlation test to examine which taxa within each dataset was associated with host age. For each dataset in which the null hypothesis of no association was true, we would expect a uniform distribution of p-values. Six of the 11 datasets appeared to have such a uniform distribution, while the other five cohorts show more of a left skew distribution of Kendall p-values, which suggest an age signature (Supp. Figure 4A).

At a 5% FDR adjusted threshold, these five cohorts (AGP, Noguiera BCN0, Baxter, Gloor, and Goodrich) had taxa significantly correlated with age. Interestingly, these five cohorts also have the larger sample sizes among the 11 datasets (Supp. Figure 4A).

### There are significant reproducible associations of age between some of the cohorts with larger sample sizes

After assessing each dataset for significant associations between host age and specific taxa, we next looked to determine how consistent microbial shifts were across datasets. We took the Kendall p-values from the prior analysis and compared all pairs of cohorts for a total of 55 pairwise comparisons (11 choose 2) of log-transformed Kendall p-values (Figure 4) visualized using p-value vs. p-value plots (Fig. 1).

**Figure 4.**
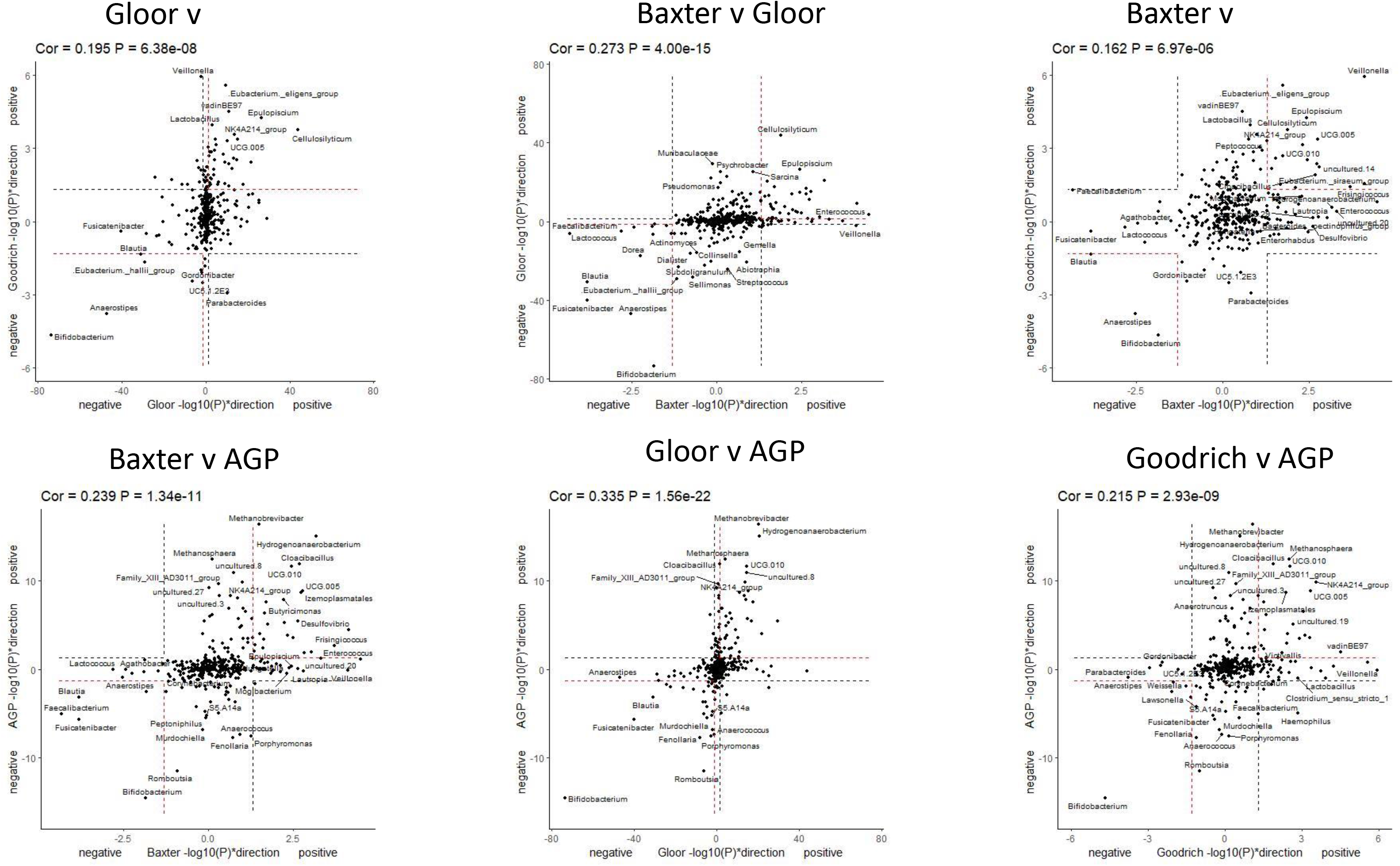
Pairwise comparisons of unadjusted Kendall p-values. (A-D) Pairwise comparisons between the four largest cohorts in this study: the AGP, Goodrich, Gloor, and Baxter cohorts. In these plots, we labeled the taxa that were significantly correlated with host age in both cohorts (seen in the lower left and upper right quadrants).

Out of the 55 pairs, we observed 21 pairs with significant Kendall correlation (p-value< 0.05). Most of the significantly correlated pairs include cohorts with larger sample sizes, particularly the Gloor, the Goodrich, Baxter, and AGP cohorts. Most datasets had no correlation, and some even had a negative correlation (Kendall tau), including the Nogueira (BCN0) and Escobar pair. We observed similar results in both parametric and non-parametric models (Supp. Figure 5).

### Power analysis suggests a large sample size of ~100-300 subjects is needed to observe an age signal in human gut microbiome data

Since the effect size of the age signal is modest, future studies will need to explicitly consider what sample size will be needed to successfully reveal these associations. To guide such work, we performed a power analysis by randomly subsampling without replacement to determine the minimum sample size required in each cohort to see significant associations. We calculated the number of significant taxa (FDR < 0.05) at the rarefied sample sizes for each cohort and observed that we can reliably detect taxa with a sample size between 100-300 samples (Figure 5). Similar results from these power estimates were observed for parametric and non-parametric tests (Supp. Figure 6 and 7).

**Figure 5.**
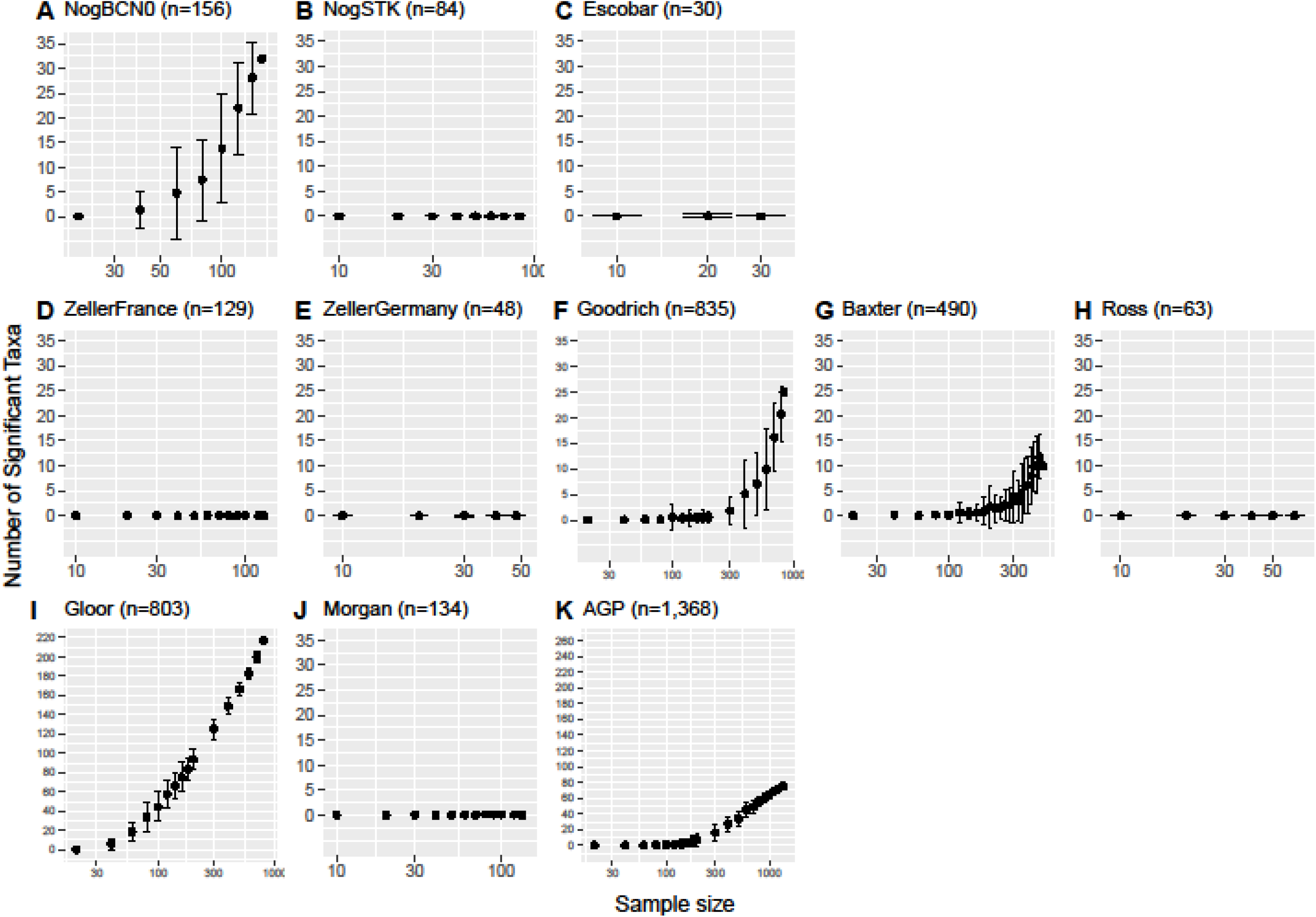
Rarefaction plots comparing the number of significant taxa at incremented sample sizes for each dataset, which are log_10_-scaled along the x-axis. We repeated the correlation test at each sample size 50 times, with each dot representing the average number of significant taxa per sample size, and the error bars represent the ± standard deviation.

### Age signature within the human gut microbiome data is not entirely driven by children or elderly individuals

In the Kendall correlation tests and power analysis, we observed taxa significantly associated with host age within the larger cohorts, but we did not see any trends that were specific to any of the three subgroups (Figure 5, Supp. Figure 6). As such, observing taxa that were significantly associated with host age was not limited to datasets with a larger age range. Rather, we observed significant taxa within cohorts with larger sample sizes.

However, to rule out the possibility that samples from extreme age ranges are driving the observed age signal, we repeated the power analysis with subsets of the AGP, Gloor, Goodrich, and Baxter cohorts that exclude the elderly population (aged 80 years and older) and subsets that exclude the younger population (aged 17 or younger) (Supp. Figure 8). Datasets excluding the elderly population and the younger population exhibited similar trends of significant taxa to the full respective datasets. Overall, we conclude that the observed age signal in these studies was not driven entirely by extreme age groups (Supp. Figure 8).

### No single taxon is consistently associated with aging across all studies

There were 65 genera (excluding those with unspecified labels such as “uncultured group” or “UCG”) that were present in all 11 cohorts. Of these 65 shared taxa, none were statistically significant at a 5% FDR threshold across all cohorts. At most, we observed eight taxa (*Bifidobacterium, Faecalibacterium, Blautia, Eubacterium._siraeum_group, Haemophilus, Bacteroides, Howardella,* and *Parabacteroides*) that were statistically significant across three cohorts.

To compare correlation between age and genus abundance across cohorts, we plotted the Pearson correlation R value for these eight taxa (Figure 6). The null hypothesis is indicated by the dashed line at zero, which postulates that there is no correlation between age and taxa. *Bifidobacterium* had negative correlations in eight cohorts, but positive correlations in the Ross, Nog_STK, and Escobar cohorts (Figure 6). To rule out count sparsity as a potential factor driving the signal, we compared the total zero counts in each dataset (Figure 6, Supp. Figure 9).

**Figure 6.**
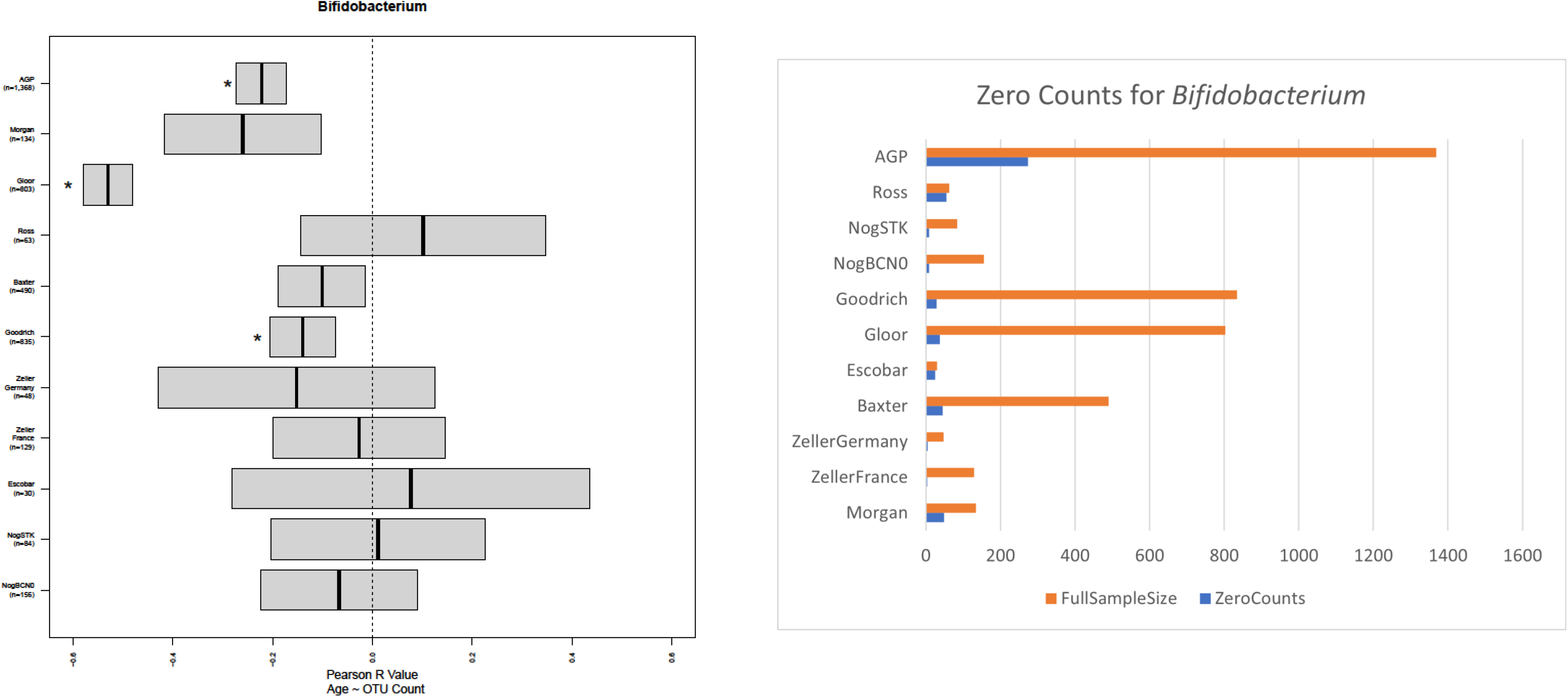
(Left) The confidence intervals for each Pearson R value, indicated by the ends of each boxplot, for *Bifidobacterium*. The dotted line at zero indicates the null hypothesis of no correlation between changes in genus abundance and host age, while the asterisks indicate the cohorts with significant correlation (Kendall p-value 5% threshold). (Right) A visualization of count sparsity by comparing the total number of samples with no counts for *Bifidobacterium* to the total sample size per cohort.

In a final analysis, we asked how many taxa present in all cohorts also shared the same correlation direction across all 11 cohorts. Among the 65 common taxa, no taxa had the same correlation direction (either all negative or all positive correlations) across all cohorts. There were two taxa that had negative correlations in ten out of 11 cohorts (*Faecalibacterium, Dialister*) and five taxa with positive correlations in ten out of 11 cohorts (*Butyricimonas, Izemoplasmatales, Colidextribacter, .Eubacterium._nodatum_group, Cloacibacillus*).

However, by a simple binomial distribution in which the null hypothesis that there are no taxa associated with host age was always true, we would expect one taxon to correlate in the same direction across all 11 cohorts. Similarly, we would still expect to see around one taxon correlate in the same direction across ten out of 11 cohorts. As such, our observations are more than what we would expect to see by chance, particularly with the positively correlated taxa. This analysis is consistent with our overall conclusion that there is not strong evidence of taxa that share a similar pattern of association with age across all 11 cohorts in our study.

### Discussion

While we observed significant age-related microbial shifts in most cohorts, there were no significant taxa in common across all 11 cohorts in our integrative analysis. We did observe eight significant taxa (*Bifidobacterium, Faecalibacterium, Blautia, Eubacterium._siraeum_group, Haemophilus, Bacteroides, Howardella,* and *Parabacteroides*) in common across three of our cohorts with the largest sample sizes. In the four largest cohorts (AGP, Goodrich, Gloor, Baxter), we observed some consistent reproducibility of the age-signal with significant correlations and modest Kendall tau values (Figure 4) in p-value vs. p-value plots. Our results are most consistent with a conceptual model of aging in which there are modest consistent associations between some cohorts in the gut microbiome (Figure 1, middle panels), but a consistent age signal was not detected across all eleven datasets. We observed reproducible age signals among the larger cohorts, which included populations from the USA, UK, and China, which indicates that support for a consistent “biological clock hypothesis” is likely to be seen across even studies from very different regions given sufficient sample sizes.

Interestingly, in the larger studies, we observed consistent results with both parametric and non-parametric models. The parametric model had slightly smaller ranges of p-values when comparing the correlation between a given pair of cohorts for the p-value comparison plots, but the overall results were similar. Our results therefore suggest that non-parametric models can successfully be used by investigators who wish to avoid the assumptions of linear regressions. Regardless of the statistical model used, our power estimates suggest a minimal sample size of approximately 100-300 samples to reliably detect an age signal in the human gut microbiome.

A significant limitation of our study is integrating data across laboratories that have different extraction and sequencing protocols. This unquestionably adds bias to each of the datasets which could obscure a common signal. However, other integrative analyses performed by our lab have successfully established robust signatures across samples collected and sequenced under different protocols (37). Moreover, integrative analyses for diseases such as Crohn’s disease and ulcerative colitis have also been successful at identifying common signals across studies (38). Therefore, it is reasonable to assert that if there were a strong signal associated with aging that was reliable across datasets and detectable with small sample sizes, then it is likely that our integrative approach would have successfully detected it.

Another limitation of our analysis is that we examined only datasets that contain a cross-sectional view of host aging and the gut microbiome, excluding longitudinal sampling from individuals over time. A study with longitudinal data would overcome individual differences in the microbiome and might provide a stronger signal of change across studies. Access to longitudinal data is often limited, and the time range of longitudinal samples varies between independent studies. Of the 11 cohorts in this analysis, four datasets had some longitudinal sampling with varying timepoints, but to maintain consistency across all cohorts, we focused on the cross-sectional data in this analysis. Our study is also limited by using only 16S rRNA gene sequence data, which limits our analysis to the genus level. Whole genome sequencing (WGS) data allows for a more refined taxonomic resolution and would also yield a functional perspective of the aging gut microbiome.

A recent study showed that both stool and rectal swab samples can give more similar inference to each other than when compared with mucosal tissue samples (39). We note that while most of the datasets in this analysis are based on stool samples, the Zeller Germany cohort consisted of only tissue samples. We did not see significant correlations with age within the Zeller Germany cohort, which was also one of the smaller cohorts (n=48). The Morgan study had both stool (n=134) and colonic tissue (n=94), but in this study we only focused on the stool, although there were no significant associations with age in either tissue type. When we combined the stool and colonic samples together to make 228 samples (data not shown), we did see a single taxon (*Bifdobacterium*) significantly negatively associated with age, giving some further support to the idea that *Bifdobacterium* is consistently associated with age in cohorts with large sample sizes and is the most reliable biomarker of age in the human associated microbiome. However, it is unclear if combining the samples in this way is statistically valid for the Morgan study, so we did not emphasize this result from the combined samples in this manuscript. It will be of great to interest to see if our overall *Bifdobacterium* results from three of the four largest datasets are replicated in future cohorts and if future studies are able to mechanistically link these changes to *Bifdobacterium* with changes to the function of the gut microbiome with advancing age.

Our study extends a previously published meta-analysis looking at age and sex differences (24). That study looked at 4 datasets, two of which were part of the American Gut Project cohort (also used within our study), and did not explicitly consider sample size. Like our study, the previous study found alpha diversity differences in some samples but not others. The machine learning approach taken by that study had some success in linking microbiome age to chronological age consistent with our arguments that age has some consistent associations with the microbiome. Our paper did not emphasize machine learning as we instead focused on which taxa were reproducible across studies and how that reproducibility was impacted by sample size. It will be of interest to see if future machine learning approaches in general become more powerful, building on the models proposed by de la Cuesta-Zuluaga et. al, as more cohorts with large sample sizes become publicly available to serve as training sets.

Our study argues that there are some consistent changes to the microbiome across a wide geographical range. While of course the associations we describe do not prove a causal mechanistic relationship, the hypotheses generated by our integrative analysis are intriguing in that they raise the possibility that understanding the relationship between these changes in the microbiome and host age could potentially provide insight for clinical applications. For example, if the changes to the microbiome cause increased disease risk or risk of medicinal side effects in older people, it might be possible to mitigate this risk by shaping the microbial community with probiotics, prebiotics, or other interventions. Future studies will undoubtedly generate additional insights into whether changing the microbiome to a profile that matches younger subjects can promote longevity and healthy aging.

## Authors’ Contributions

ADM: Data curation, formal analysis, investigation, visualization, manuscript: writing original draft, review and edit; SS: Methodology, visualization, testing, manuscript: review and edit; AAF: Conceptual development, supervision, methodology, visualization, manuscript: writing original draft, review and edit; ICB: Testing, manuscript: review and edit.

## Availability of data and materials

The datasets and metadata in this analysis are publicly available in public databases, with accession numbers in Table 1 and 2. Scripts used in this analysis are available at https://github.com/anhmoss/AgingMicrobiome_IntegrativeAnalysis.

## Ethics approval and consent to participate

Not applicable.

## Consent for publication

Not applicable.

