## Supplemental Figures for "An integrative analysis of the biological clock hypothesis in human gut microbiome"

Anh Moss

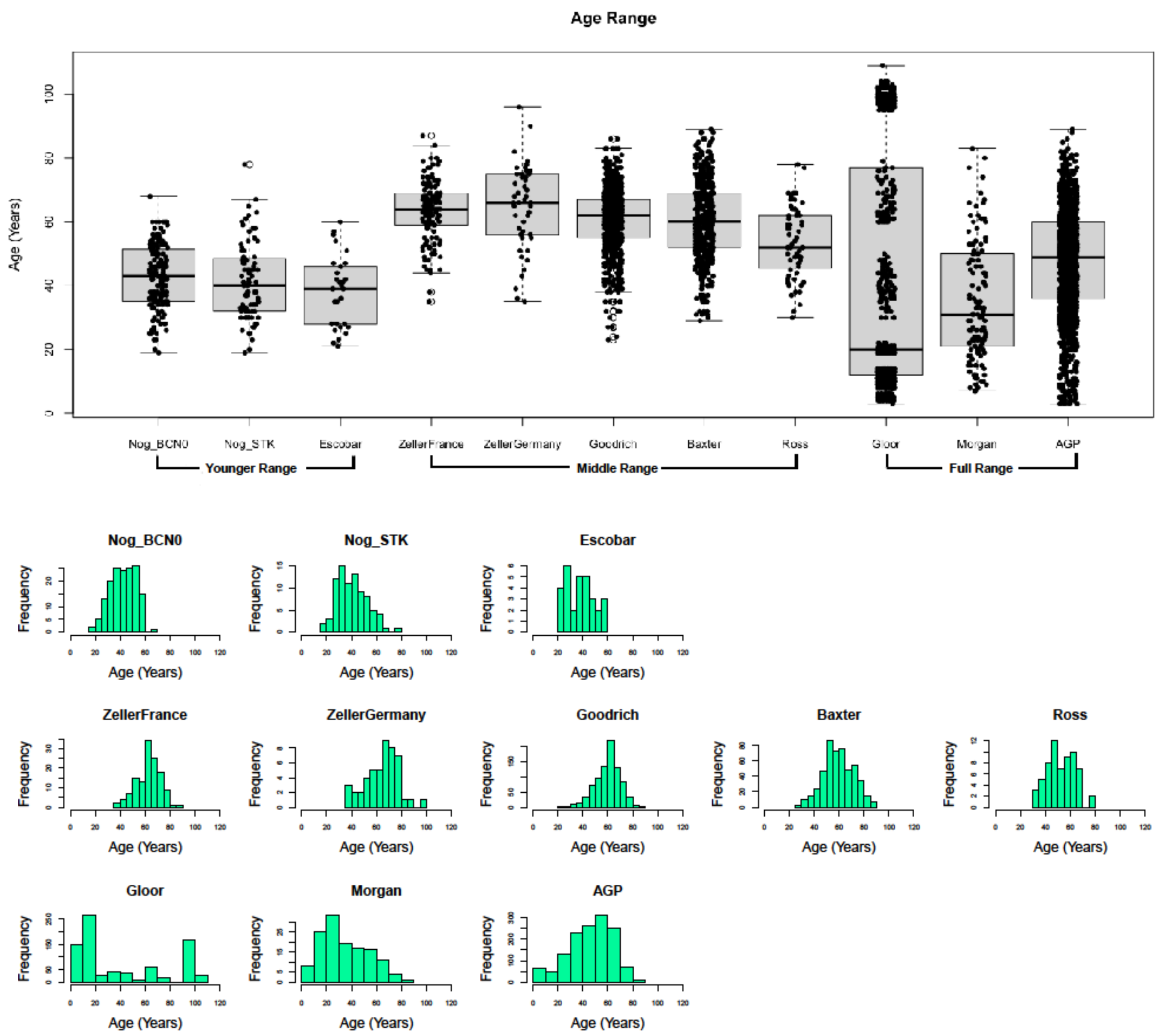


**Supp. Figure 1**. Visual summaries of the age ranges for each of the 11 datasets.

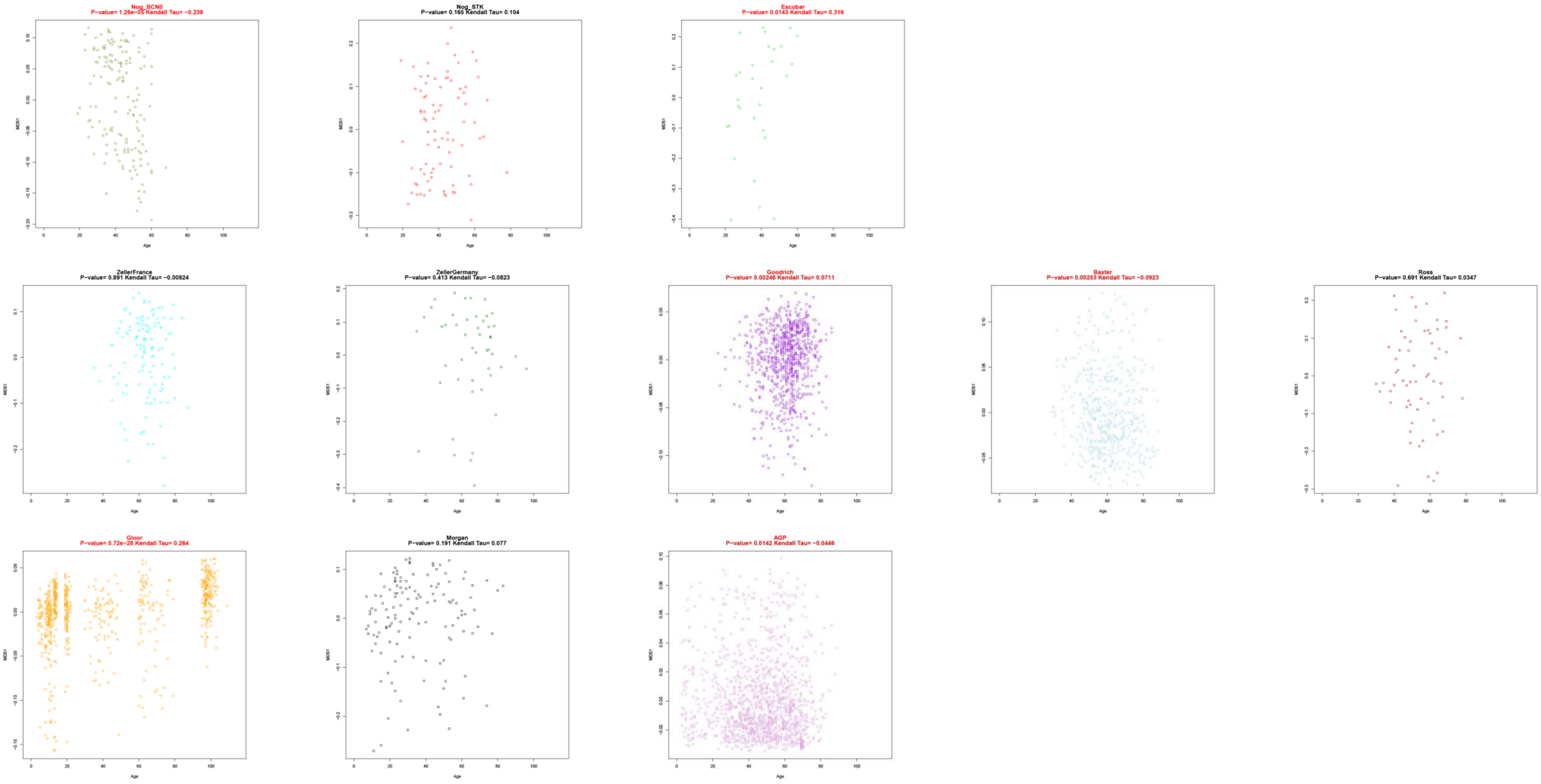


**Supp. Figure 2A.** Scatterplots of age regressed against MDS1, with Kendall tau and p-values.

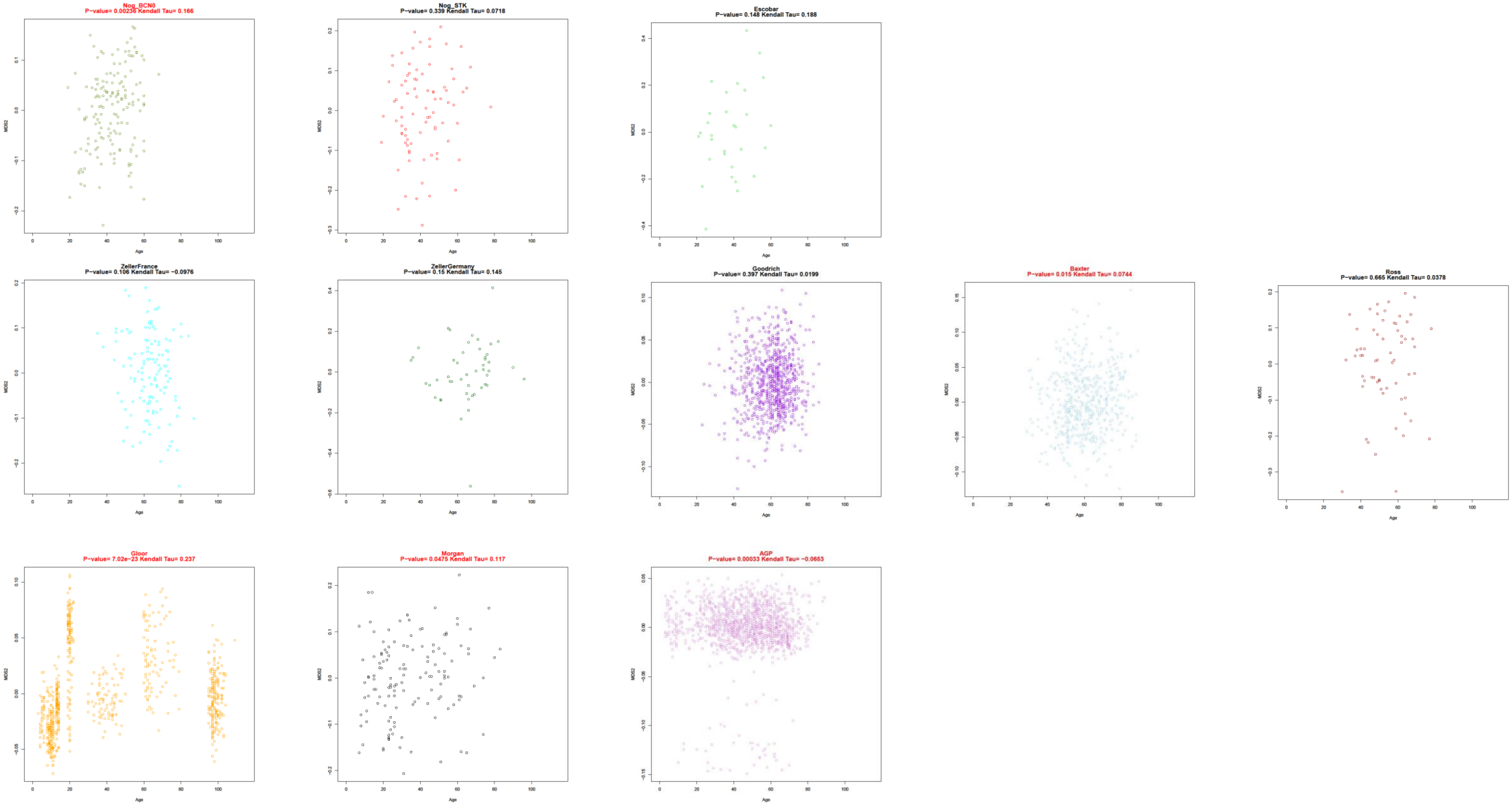


**Supp. Figure 2B.** Scatterplots of age regressed against MDS2, with Kendall tau and p-values.

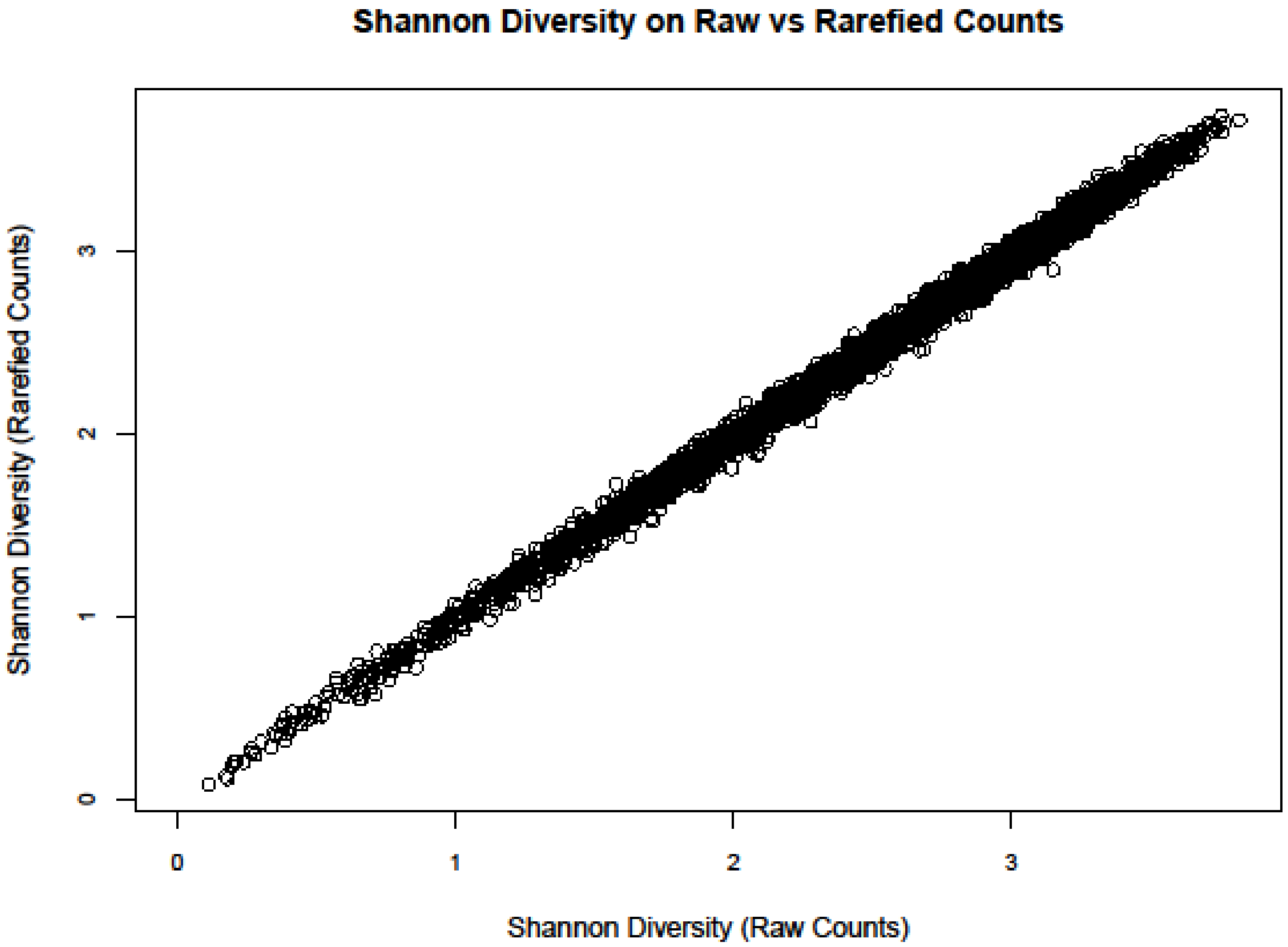


**Supp. Figure 3**. Comparing Shannon Diversity with raw counts and rarefied counts (threshold with 1000 counts). Results for Shannon diversity were similar between raw and rarefied counts.

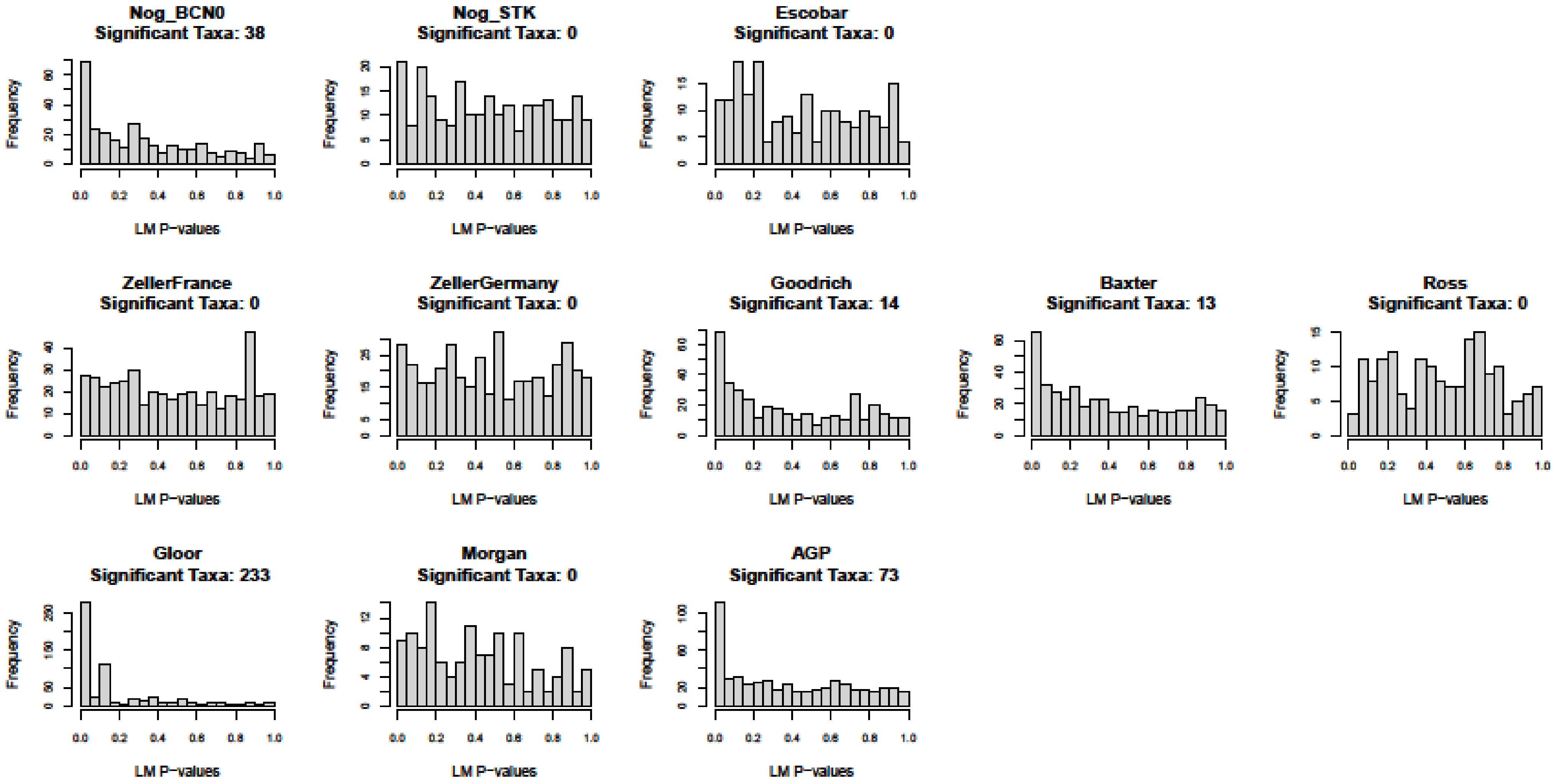


**Supp. Figure 4A.** Histogram of Kendall p-values for each cohort. Significant taxa refers to taxa with abundance shifts significantly associated with host age (FDR 5% threshold).

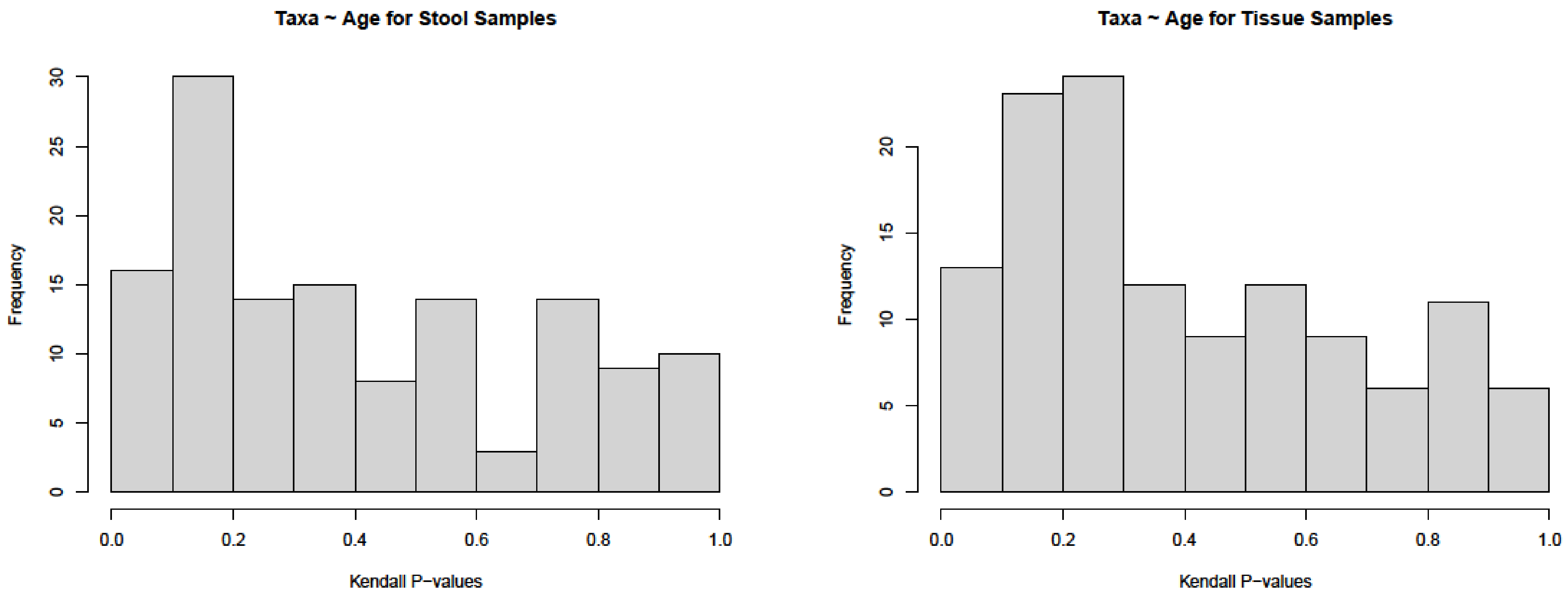


**Supp. Figure 4B.** Kendall p-value histograms for stool and tissue samples in the Morgan cohort.

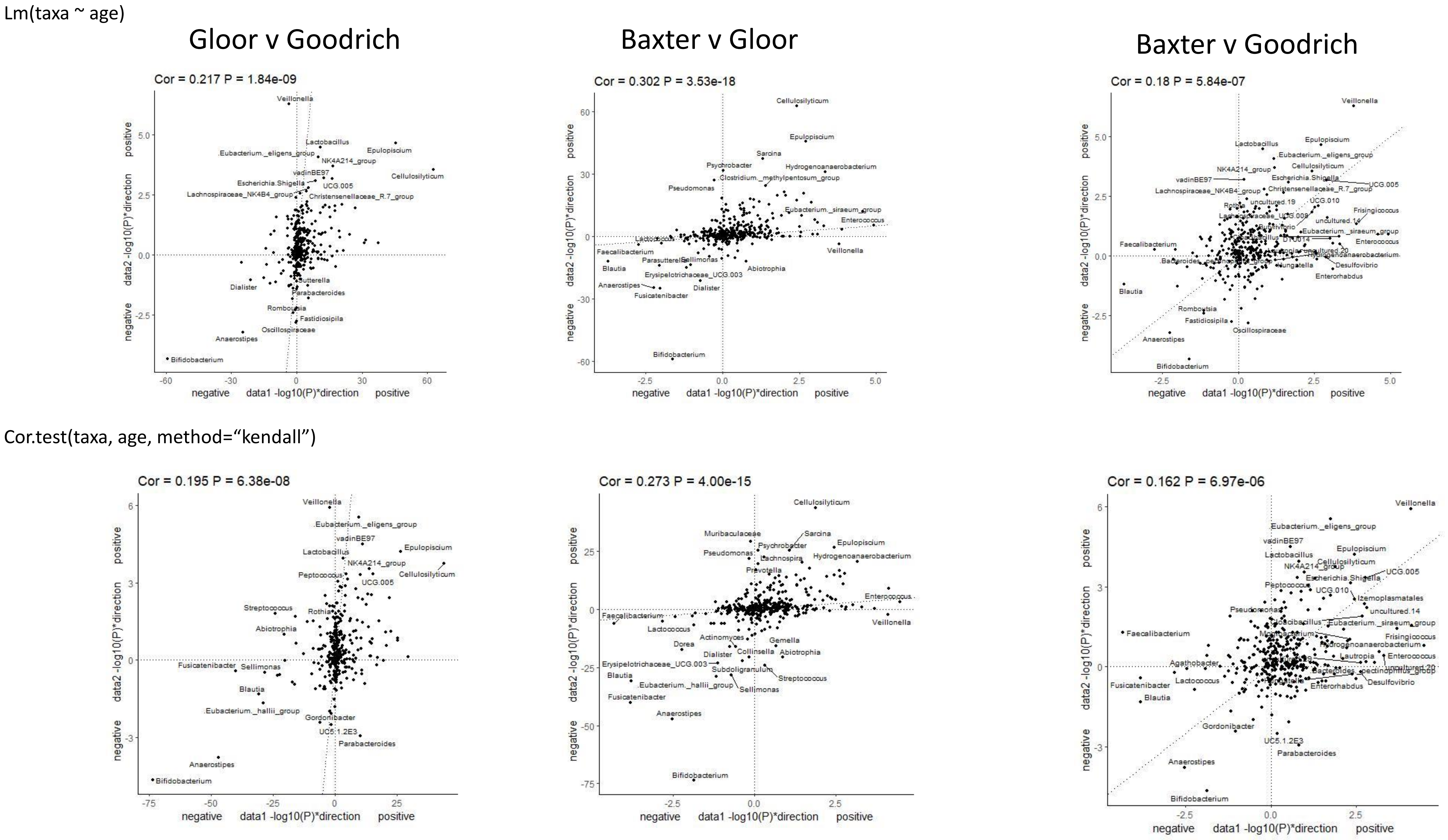


**Supp. Figure 5A**. P-value vs. p-value comparisons with parametric (simple linear regression, top) and non-parametric(Kendall correlation, bottom) models for the pairwise comparisons of the 4 largest datasets in our analysis.

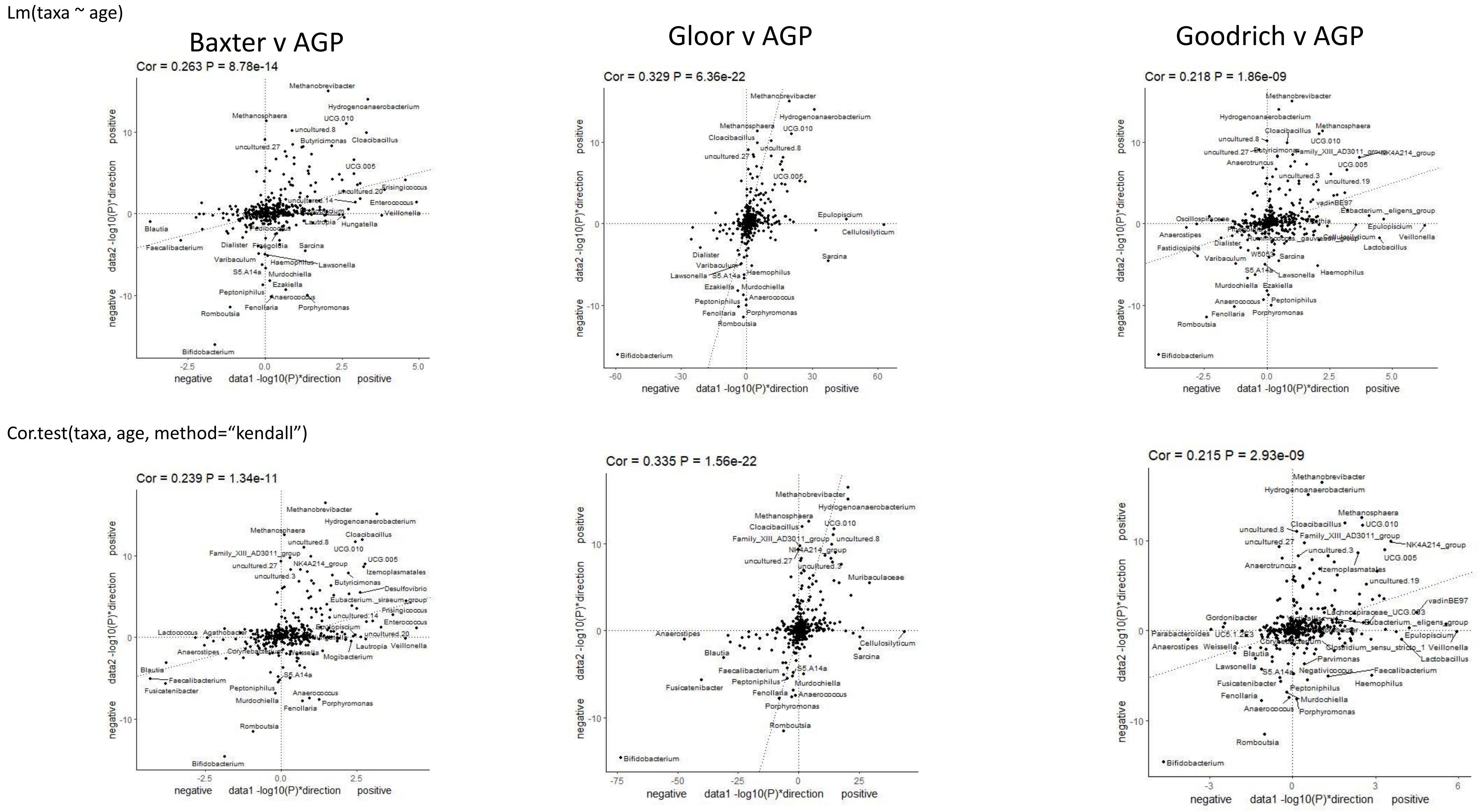


**Supp. Figure 5B**. P-value vs. p-value comparisons with parametric (simple linear regression, top) and non-parametric(Kendall correlation, bottom) models for the pairwise comparisons of the 4 largest datasets in our analysis.

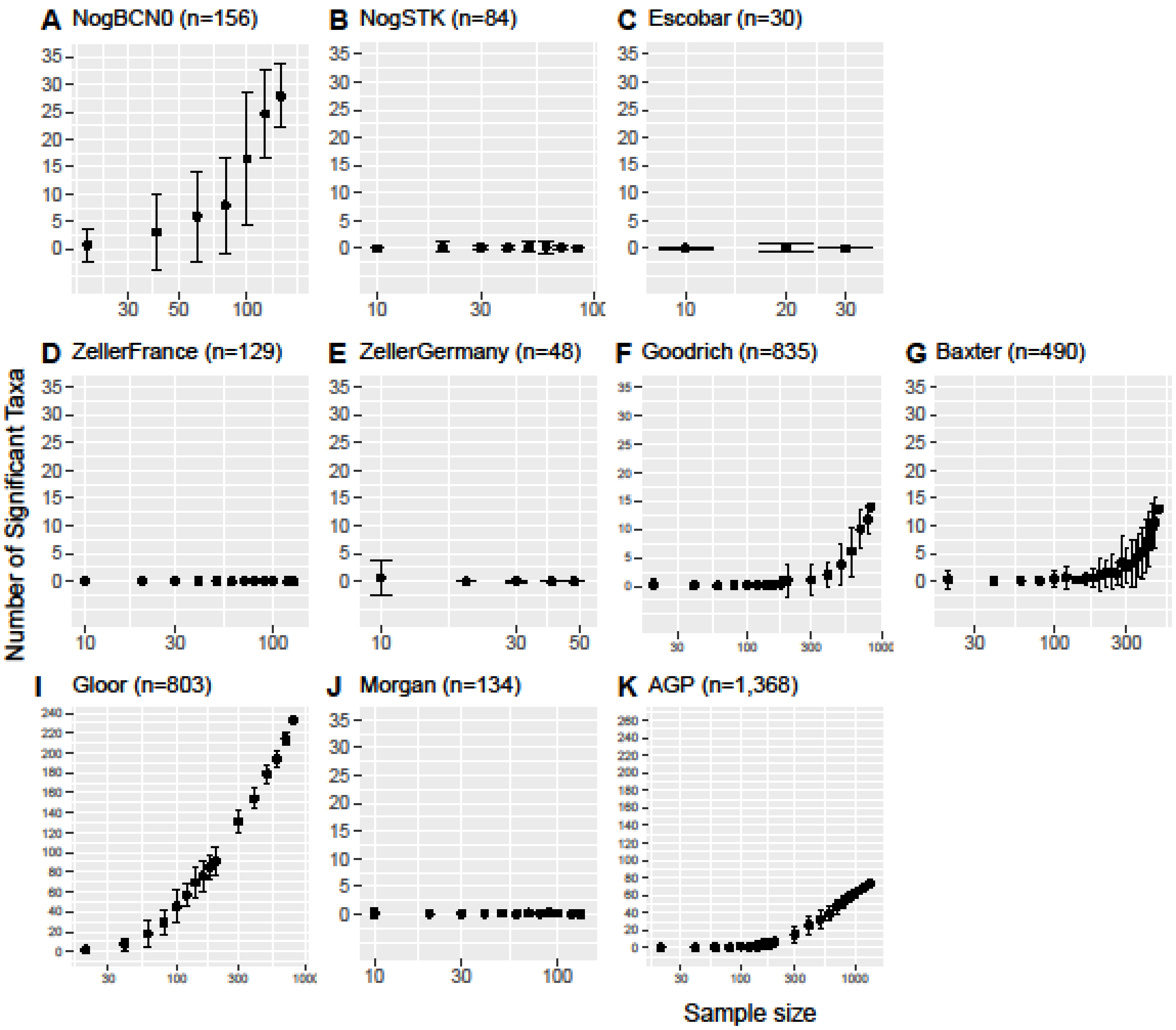


**Supp. Figure 6.** Power analysis with parametric modeling (simple linear regression) for each cohort.

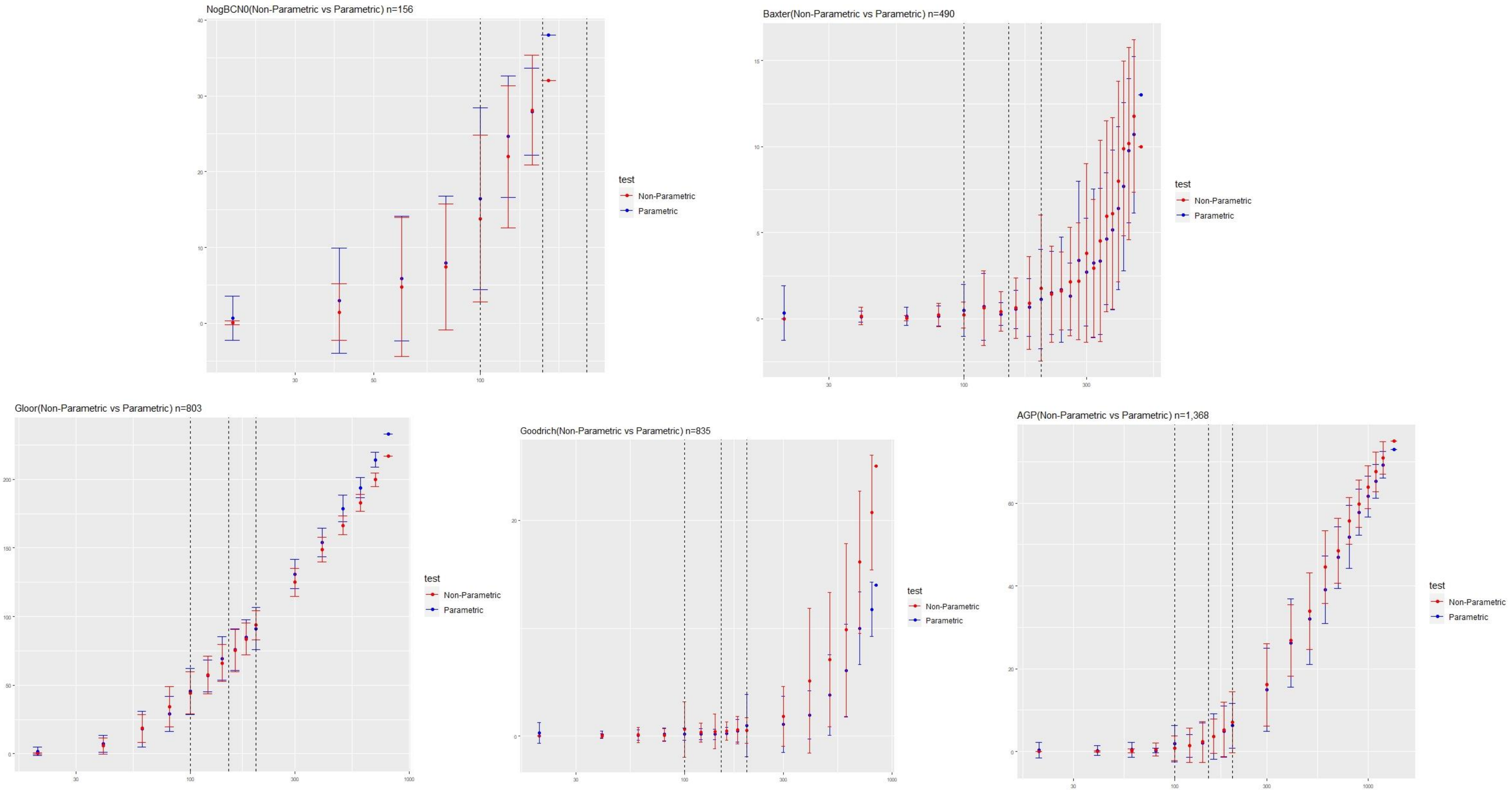


**Supp. Figure 7.** Close ups on power analysis plots, comparing both nonparametric (red) and parametric (blue) results. Five out of 11 cohorts had taxa significantly associated with host age at a 5% FDR threshold, hence only five plots are presented.

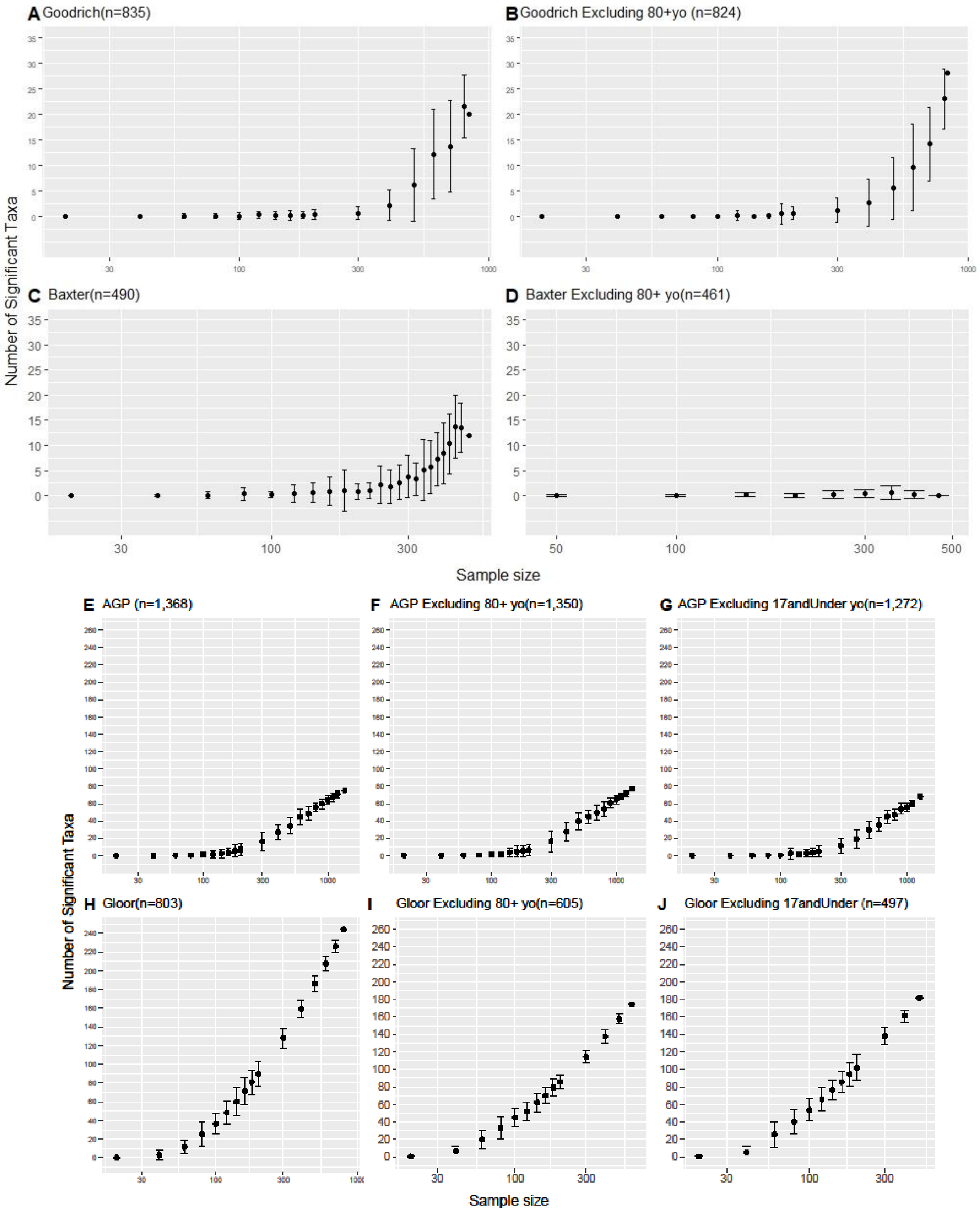


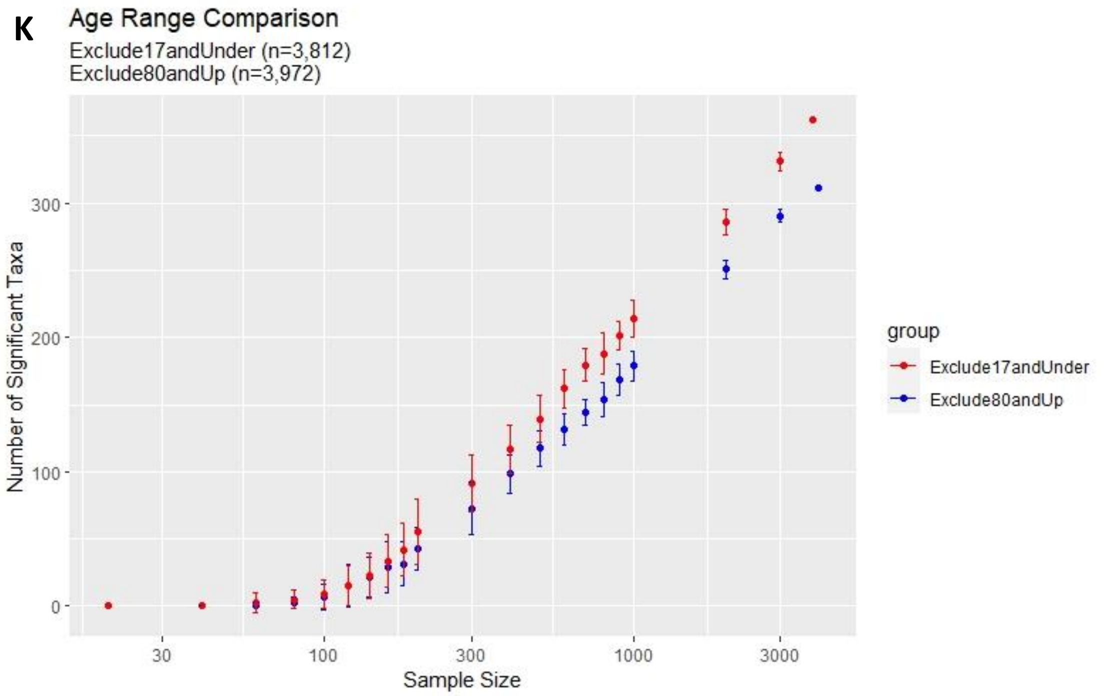


**Supp. Figure 8.** We compare the impact of our power analysis with two ranges of host ages, children (ages 3 to 17 years) and elderly (ages 80 and up). In these power rarefaction plots, we compare the age signal with the full dataset and with samples excluding either children and/or elderly participants (figures A-J). (K) We also pooled all samples from across the 11 cohorts to compare the collective sample without child participants (red) and with a group that excluded the elderly participants (blue). We observed that regardless of excluding samples from older or younger individuals, the age signal remains similar.

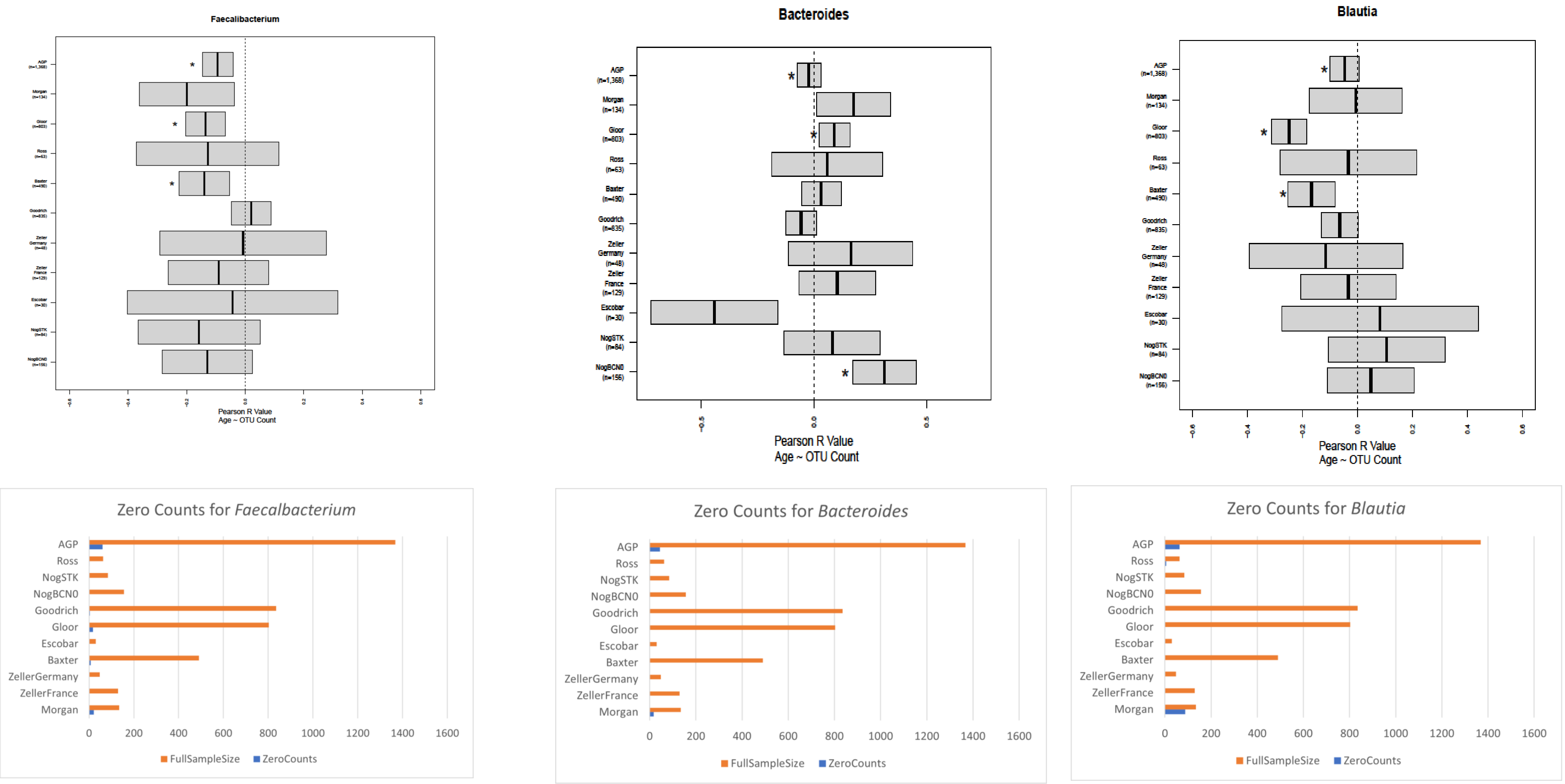


**Supp. Figure 9**. (Top) Boxplots that visualize the confidence intervals for Pearson R value (Age ~ OTU) for the other seven taxa that were significant across three different cohorts: *Faecalibacterium*, *Blautia*, *Eubacterium._siraeum_group*, *Haemophilus*, *Bacteroides*, *Howardella*, and *Parabacteroides.* Asterisks indicate the cohorts with significant correlation (Kendall p-value 5% threshold). (Bottom) A visualization of count sparsity by comparing the total number of samples with no counts for *Bifidobacterium* to the total sample size per cohort.

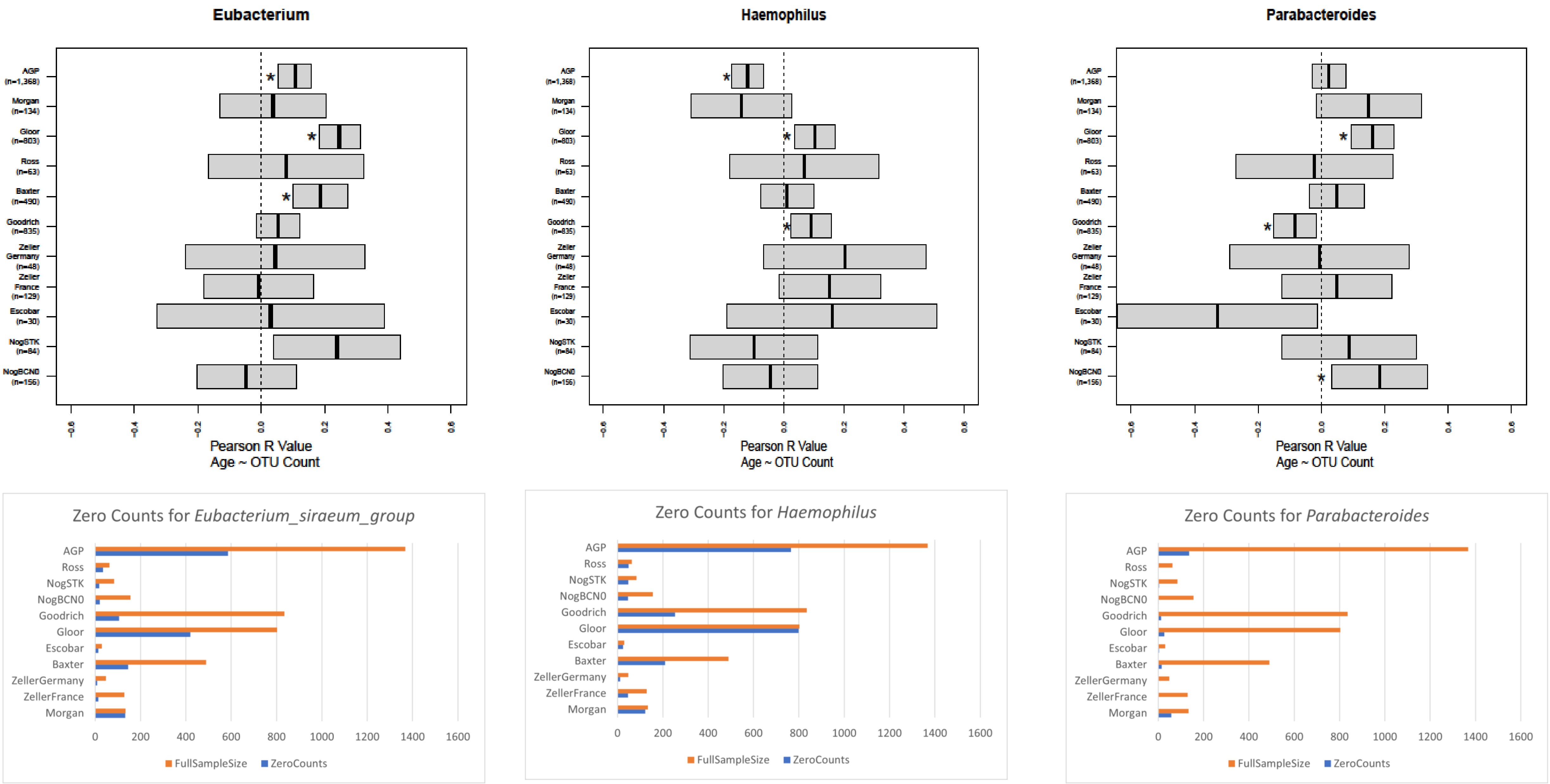


**Supp. Figure 9**. (Top) Boxplots that visualize the confidence intervals for Pearson R value (Age ~ OTU) for the other seven taxa that were significant across three different cohorts: *Faecalibacterium*, *Blautia*, *Eubacterium._siraeum_group*, *Haemophilus*, *Bacteroides*, *Howardella*, and *Parabacteroides.* Asterisks indicate the cohorts with significant correlation (Kendall p-value 5% threshold). (Bottom) A visualization of count sparsity by comparing the total number of samples with no counts for *Bifidobacterium* to the total sample size per cohort.

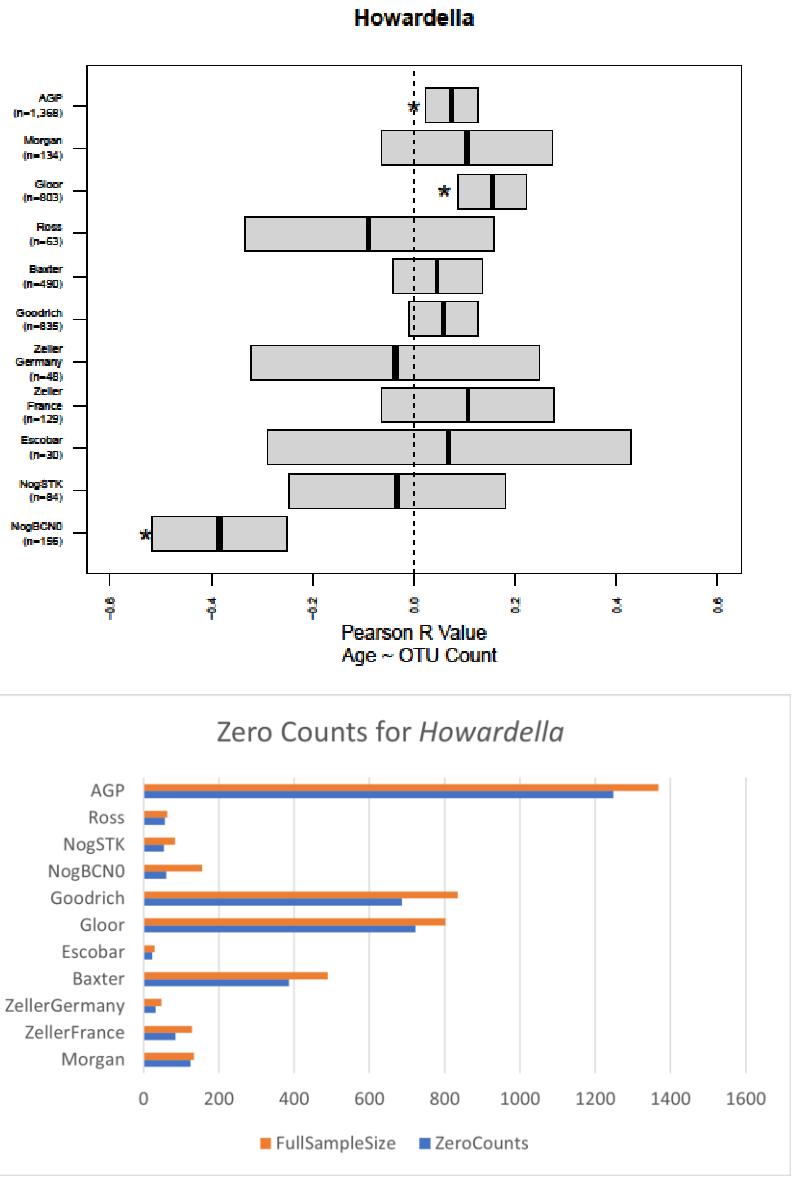


**Supp. Figure 9**. (Top) Boxplots that visualize the confidence intervals for Pearson R value (Age ~ OTU) for the other seven taxa that were significant across three different cohorts: *Faecalibacterium*, *Blautia*, *Eubacterium._siraeum_group*, *Haemophilus*, *Bacteroides*, *Howardella*, and *Parabacteroides.* Asterisks indicate the cohorts with significant correlation (Kendall p-value 5% threshold). (Bottom) A visualization of count sparsity by comparing the total number of samples with no counts for *Bifidobacterium* to the total sample size per cohort.
